# The general transcription factor TFIIB is a target for transcriptome control during cellular stress and viral infection

**DOI:** 10.1101/2024.01.16.575933

**Authors:** Leah Gulyas, Britt A. Glaunsinger

**Affiliations:** Department of Plant and Microbial Biology, University of California, Berkeley, CA 94720, USA; Department of Molecular and Cell Biology, University of California, Berkeley, CA 94720, USA; Howard Hughes Medical Institute, University of California, Berkeley, CA 94709, USA

**Keywords:** TFIIB, general transcription factor, RNAPII transcription, KSHV, caspase-3 cleavage, proteasomal turnover

## Abstract

Many stressors, including viral infection, induce a widespread suppression of cellular RNA polymerase II (RNAPII) transcription, yet the mechanisms underlying transcriptional repression are not well understood. Here we find that a crucial component of the RNA polymerase II holoenzyme, general transcription factor IIB (TFIIB), is targeted for post-translational turnover by two pathways, each of which contribute to its depletion during stress. Upon DNA damage, translational stress, apoptosis, or replication of the oncogenic Kaposi’s sarcoma-associated herpesvirus (KSHV), TFIIB is cleaved by activated caspase-3, leading to preferential downregulation of pro-survival genes. TFIIB is further targeted for rapid proteasome-mediated turnover by the E3 ubiquitin ligase TRIM28. KSHV counteracts proteasome-mediated turnover of TFIIB, thereby preserving a sufficient pool of TFIIB for transcription of viral genes. Thus, TFIIB may be a lynchpin for transcriptional outcomes during stress and a key target for nuclear replicating DNA viruses that rely on host transcriptional machinery.

**Graphical Abstract:** 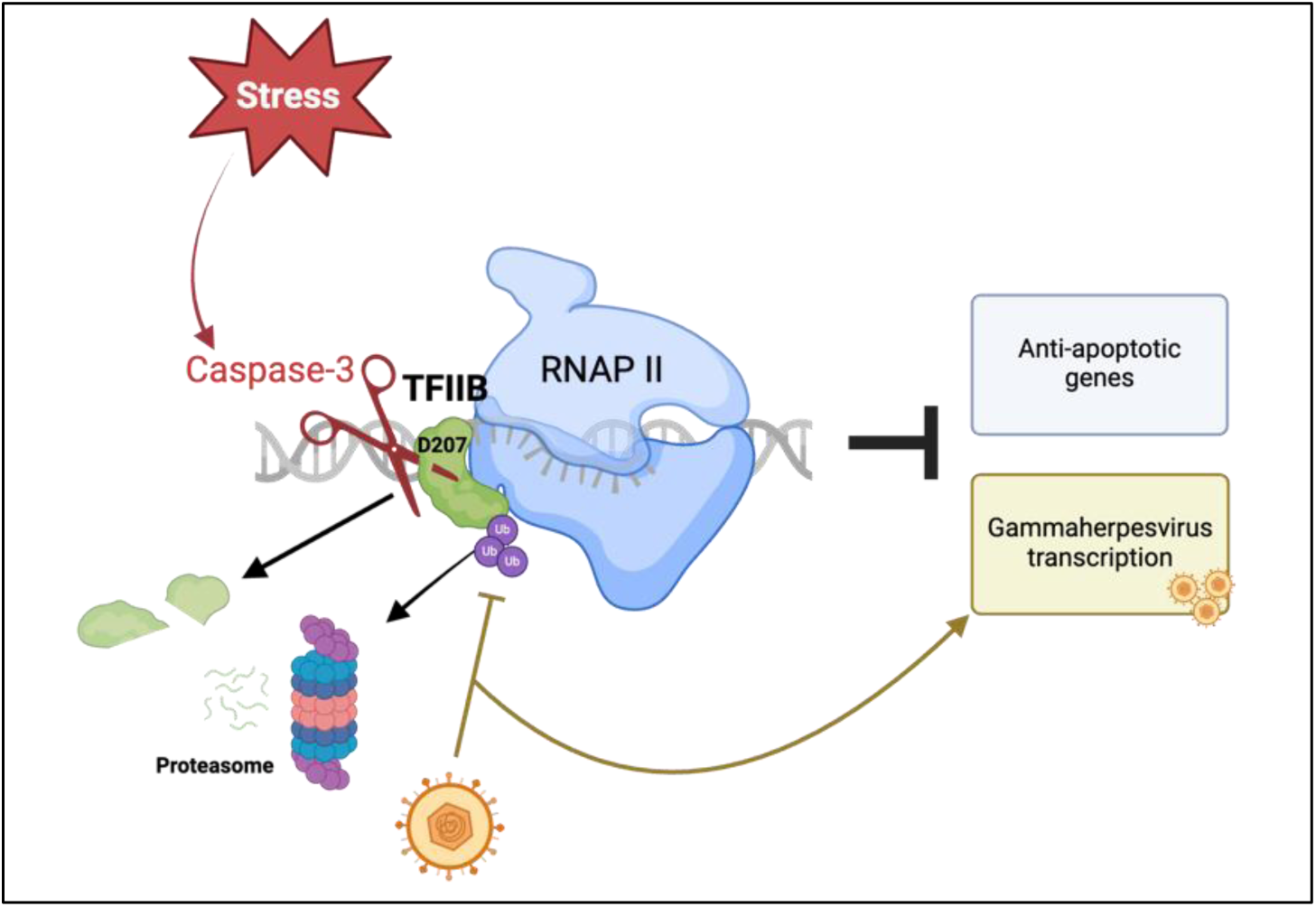

**Significance Statement:** Transcription by RNA polymerase II (RNAPII) synthesizes all cellular protein-coding mRNA. Many cellular stressors and viral infections dampen RNAPII activity, though the processes underlying this are not fully understood. Here we describe a two-pronged degradation strategy by which cells respond to stress by depleting the abundance of the key RNAPII general transcription factor, TFIIB. We further demonstrate that an oncogenic human gammaherpesvirus antagonizes this process, retaining enough TFIIB to support its own robust viral transcription. Thus, modulation of RNAPII machinery plays a crucial role in dictating the outcome of cellular perturbation.

## Introduction

Cell stress is accompanied with rapid changes to the transcriptome. This involves the induction of genes required for stress resolution (1), for example through post-translational regulation of stress-responsive transcription factors like p53 and Rb (2, 3). Conversely, most stressors, including DNA damage (4), pro-apoptotic signals (5), hyperosmotic stress (6), and some viral infections, broadly downregulate basal gene expression by suppressing RNA polymerase II (RNAPII) activity (7–10). This may focus gene expression on the stress response by freeing resources for transcription and translation and may also restrict viral access to essential transcription machinery akin to the well documented translational shutdown pathways (1, 11–13). However, the mechanisms underlying this transcriptional repression remain unclear.

DNA viruses are valuable models for understanding stress response coordination, as they induce cell stress yet also rely on core cellular gene expression pathways that are frequently inactivated during stress. One DNA virus that induces a broad shutdown of host gene expression is Kaposi’s sarcoma-associated herpesvirus (KSHV), an oncogenic gammaherpesvirus that is the most common cause of cancer in individuals with HIV-associated immunosuppression. The KSHV dsDNA genome resides as a largely transcriptionally quiescent episome that is tethered to host chromatin in latently infected cells. Viral amplification and full viral gene expression occur only upon reactivation into the lytic cycle. During lytic KSHV replication, the virus usurps RNAPII for viral transcription, while dampening cellular gene expression both in the cytoplasm via accelerated mRNA decay by a viral endonuclease and in the nucleus via suppression of RNAPII transcription (7, 9, 14, 15). The cellular transcriptional repression is likely a consequence of multiple virus-induced phenotypes, including a nuclear influx of RNA binding proteins as well as altered levels of core transcription proteins (7–9, 16). Among these, a recent proteomics study identified general transcription factor IIB (TFIIB, also known as GTF2B) as one of the most prominently depleted transcription factors in gammaherpesvirus infected cells (7).

TFIIB is a core member of the RNAPII holoenzyme and is crucial for RNAPII recruitment and initiation (17). It is a small, ∼35 kD protein that stabilizes the TBP-containing general transcription factor IID (TFIID) on promoter DNA near the TFIIB C-terminus (17, 18). TFIIB serves as bridge between TFIID and RNAPII, with its N-terminal B-ribbon portion threading to the RNAPII core complex and drawing it to the promoter. Once bound, TFIIB assists RNAPII in unwinding DNA and identifying the transcription start site (18, 19). TFIIB also has a more enigmatic role in 3′ transcript processing in yeast, involving gene looping via interactions with the 3′ CPF complex and other termination machinery (20–22), as well as proper transcription termination in mammalian cells (23). Acute PROTAC-driven loss of TFIIB protein has recently been shown to have a broadly repressive effect on RNAPII activity (23).

TFIIB-dependent transcription is influenced by its phosphorylation, which is reduced during DNA damage (24, 25). If and how TFIIB is otherwise regulated remains unknown. Here, we show that TFIIB protein abundance is tightly controlled in both stressed and unstressed cells by two distinct pathways. During DNA damage, apoptosis, and infection-induced stress, caspase-3 is activated and TFIIB undergoes rapid caspase-mediated cleavage. This caspase-mediated TFIIB depletion suppresses RNAPII transcription, most prominently of pro-survival factors in cells undergoing apoptosis. However, as is characteristic of tightly regulated proteins, TFIIB is short-lived even in unstressed cells. The E3 ligase TRIM28 targets TFIIB for rapid turnover by the ubiquitin proteasome pathway, keeping its half-life short both in the absence and presence of stress. Finally, we demonstrate that KSHV, which requires RNAPII for transcription, counteracts loss of TFIIB by impeding its proteasomal degradation. This work highlights how posttranscriptional control of a core general transcription factor can influence the cellular transcriptional response to stress and can be manipulated for viral benefit.

## Results

### Cellular stress depletes TFIIB via caspase cleavage, suppressing anti-apoptotic genes

To evaluate how KSHV influences TFIIB protein levels, we used a B cell lymphoma cell line that is stably latently infected with KSHV (TRExRTA BCBL-1, henceforth referred to as BCBL-1). These cells contain an inducible version of the viral lytic activator RTA, enabling viral reactivation into the lytic cycle upon treatment with doxycycline (dox) (26).

Western blot of endogenous TFIIB protein showed that its levels declined upon lytic KSHV reactivation (**Fig. 1A)**, in agreement with prior observations with the related murine gammaherpesvirus MHV68 (7). To determine whether this depletion was specific to lytic KSHV reactivation or is also induced by other types of stress, we treated the latent cells with etoposide to cause DNA damage or cycloheximide to inhibit translation. Both treatments also downregulated TFIIB, suggesting that it may be commonly targeted during stress **(Fig. 1A)**.

**Figure 1.**
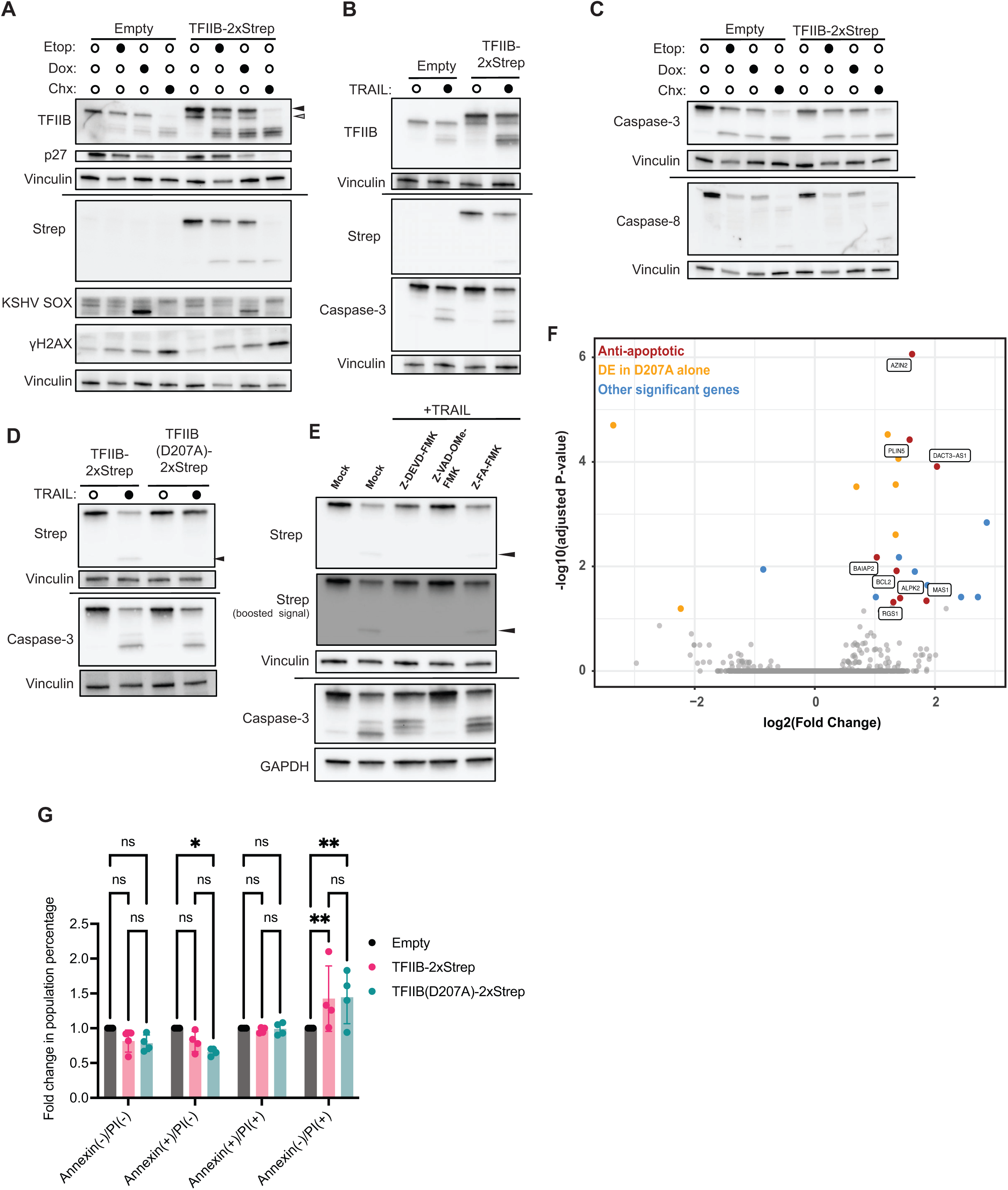
Cellular stress depletes TFIIB via caspase cleavage at D207, suppressing anti-apoptotic genes. **(A)** TRExRTA BCBL-1 cells bearing the indicated constructs were treated with 25 μM etoposide (Etop; DNA damage agent) for 24 h, 2 μg/mL doxycycline (Dox; viral reactivation) for 24h, or 100 μg/mL cycloheximide (Chx; translational inhibitor) for 6 h, or mock treated with drug diluent and analyzed by western blotting. γH2AX is induced during DNA damage response, KSHV SOX is a viral early protein and marker for infection, and p27 is a short-lived control for translation inhibition. Vinculin is a loading control. Black arrow indicates full length strep-tagged TFIIB and gray arrow indicated full-length endogenous TFIIB. **(B)** Empty or TFIIB-2xStrep expressing TRExRTA BCBL-1 cells were treated with either 100 ng/mL TRAIL ligand or mock treated with TRAIL dilution buffer for 6 h before western blotting. Caspase-3 cleavage is a control for induction of cell death. **(C)** Samples in (*A*) were probed for Caspase-3 and Caspase-8 activation by cleavage. **(D)** TRExRTA BCBL-1 cells expressing either TFIIB-2xStrep or mutant TFIIB(D207A)-2xStrep were treated with 100 ng/mL TRAIL ligand or mock for 8 h, followed by western blotting. Black arrow indicates cleavage product. **(E)** TRExRTA BCBL-1 cells expressing TFIIB-2xStrep or mutant TFIIB(D207A)-2xStrep were pretreated with 80 μM of either the caspase-3 inhibitor Z-VAD-DEVD-FMK, the pan-caspase inhibitor Z-VAD-OMe-FMK, or a negative control cysteine protease inhibitor Z-FA-FMK for 2 h. Cells were then treated with either 500 ng/mL TRAIL ligand or mock treated with TRAIL dilution buffer for 7 h followed by western blotting. Boosted strep signal has gamma contrast enhanced for visualization of the cleavage product and GAPDH probed with mouse antibody is a loading control. **(F)** TFIIB-2xStrep or TFIIB(D207A)-2xStrep expressing TRExRTA BCBL-1 cells were treated with 660 ng/mL TRAIL for 7 h and RNA extracted and sequenced with ribodepletion and ERCC spike-in control RNA for normalization. Volcano plot shows differentially expressed genes in cells containing TFIIB(D207A)-2xStrep relative to cells containing wild-type TFIIB-2xStrep. Genes in red are significantly differentially regulated in TRAIL-treated cells with described links to apoptosis (see ***SI Appendix*, Table S1**), genes in yellow are differentially expressed in both TRAIL-treated and untreated cells, and in blue are genes with unknown function or no known link to apoptosis that are differentially expressed during TRAIL-treatment. **(G)** Flow cytometry analysis of TRExRTA BCBL-1 cells treated with 25 μM etoposide for 24 h and then labelled with Annexin V-FITC and propidium iodide to assess apoptosis and permeability. Percentages of each population for TFIIB-2xStrep and TFIIB(D207A)-2xStrep cells were normalized to empty vector control. ns-nonsignificant, ** P_TFIIB(D207A)_ = 0.0061, **P_TFIIB_ = 0.0091, *P = 0.0424; two-way ANOVA with Tukey correction.

The depletion of TFIIB under all three stress conditions was associated with the appearance of smaller molecular weight bands in the ∼20 kD range. A stably introduced version of TFIIB fused to a 2x C-terminal strep tag was similarly cleaved upon treatment with etoposide, dox, and cycloheximide, suggesting that cleavage is proximal to the C-terminus of the protein (**Fig. 1A, black arrow, compared to gray arrow for the endogenous protein**). Note that p27 serves as a positive control for translation inhibition, KSHV SOX is an early protein used as a marker for viral reactivation, and γH2AX indicates the level of DNA damage. TFIIB transcriptional competence is associated with phosphorylation at residue S65 (25). However, cleavage does not depend on this phosphorylation, as both unphosphorylatable S65A and phosphomimetic S65E versions of TFIIB displayed the same degradation pattern (***SI Appendix*, Fig. S1A).**

Stress-induced proteolytic cleavage is a hallmark of caspase activity. Indeed, treatment with the TRAIL ligand to induce apoptosis caused the same TFIIB cleavage pattern (**Fig. 1B**). Furthermore, each of the cellular stressors activated both initiator caspase-8 and executioner caspase-3, as measured by a shift from inactive procaspases to their cleaved, active forms (**Fig. 1C**). Using the CaspSite database, an experimental repository for caspase substrates (27), we found a putative caspase-3 cut site after TFIIB aspartic acid residue 207 (D207). Mutation of this site to alanine (D207A) in the strep-tagged TFIIB rendered the protein resistant to cleavage, confirming this as the primary cleavage site **(Fig. 1D)**. Furthermore, both the pan-caspase inhibitor Z-VAD-OMe-FMK and the caspase-3 inhibitor Z-DEVD-FMK impaired TFIIB-2xStrep cleavage during TRAIL-induced apoptosis, whereas the Z-FA-FMK negative control for caspase inhibition did not (**Fig. 1E**). Similarly, siRNA-mediated depletion of caspase-3 decreased TFIIB cleavage during stress (***SI Appendix*, Fig. S1B**).

To assess the functional role of TFIIB cleavage, we next measured gene expression in BCBL-1 cells containing wild-type TFIIB-2xStrep or TFIIB(D207A)-2xStrep upon either mock buffer or apoptosis induction by TRAIL treatment using RNA-seq (**Fig. 1F)**. Because endogenous TFIIB was still present in these cells, we anticipated this approach would identify only the genes that were most sensitive to changes in TFIIB levels. Differential gene expression (DGE) analysis identified 22 genes that differed significantly between wild-type and D207A mutant overexpression, 16 of which were specifically upregulated in the TRAIL treated cells. Notably, half of these genes have reported anti-apoptotic roles, including BCL2 (***SI Appendix*, S1 Table)** (28–37). We therefore measured how expression of TFIIB-2xStrep or TFIIB(D207A)-2xStrep influenced the frequency of apoptosis and necrosis after 24 hours of etoposide treatment using Annexin V and propidium iodide (PI) staining (38), respectively (**Fig. 1G**). While there was an equivalent percentage of cells undergoing either form of cell death (i.e., Annexin V+/PI+) in each of the samples, modest but significant differences emerged when we separated these into populations undergoing either apoptosis (Annexin V+/PI-) or necrosis (Annexin V-/PI+). Expression of TFIIB(D207A)-2xStrep caused reduced levels of apoptotic cells, in agreement with the increased expression of anti-apoptotic genes, and a compensatory increase in necrotic cells compared to empty vector (**Fig. 1G**). Cells overexpressing caspase cleavable wild-type TFIIB-2xStrep displayed an intermediate phenotype, as expected. Collectively, these data suggest that some pro-survival/anti-apoptotic genes are particularly sensitive to TFIIB depletion and cleavage of TFIIB by caspases facilitates apoptotic cell death.

### TFIIB protein is rapidly degraded by the ubiquitin-proteasome pathway in both stressed and unstressed cells

We noted that cycloheximide treatment to block translation induced more pronounced depletion of full-length TFIIB compared to the other stresses (see **Fig. 1A; *SI Appendix*, Fig. S2A**). Given that tightly regulated proteins often have short half-lives, we hypothesized that TFIIB abundance might be controlled by an additional, caspase-independent mechanism in these cells.

We first tested this by measuring the half-life of the caspase-resistant TFIIB(D207A) mutant using a pulse-chase experiment in unreactivated cells. BCBL-1 cells expressing TFIIB(D207A)- 2x strep were pulsed with the methionine analog L-Homopropargylglycine (L-HPG) to label newly synthesized protein, then chased with media lacking L-HPG. At the indicated time points, we measured TFIIB(D207A) abundance by purifying it over strep beads and performing Click chemistry to fluorescently label the L-HPG-containing protein (**Fig. 2A**). The half-life of caspase-resistant TFIIB was less than 4.5 hours (**Fig. 2A**). We observed similarly rapid turnover in a cycloheximide time course of TFIIB(D207A)-expressing BCBL-1 cells (**Fig. 2B**) and of endogenous TFIIB in a renal carcinoma cell line, iSLK, that is also latently infected with KSHV (**Fig. 2C**). iSLK cells lack the ability to robustly activate caspases and thus do not appreciably cleave TFIIB upon lytic reactivation (***SI Appendix,* Fig. S2B**) or cycloheximide treatment **(*SI Appendix*, Fig. S2C)**. Together, these results indicate that TFIIB is subject to rapid basal turnover, consistent with a previous report in cycloheximide-treated HEK293T cells (39).

**Figure 2.**
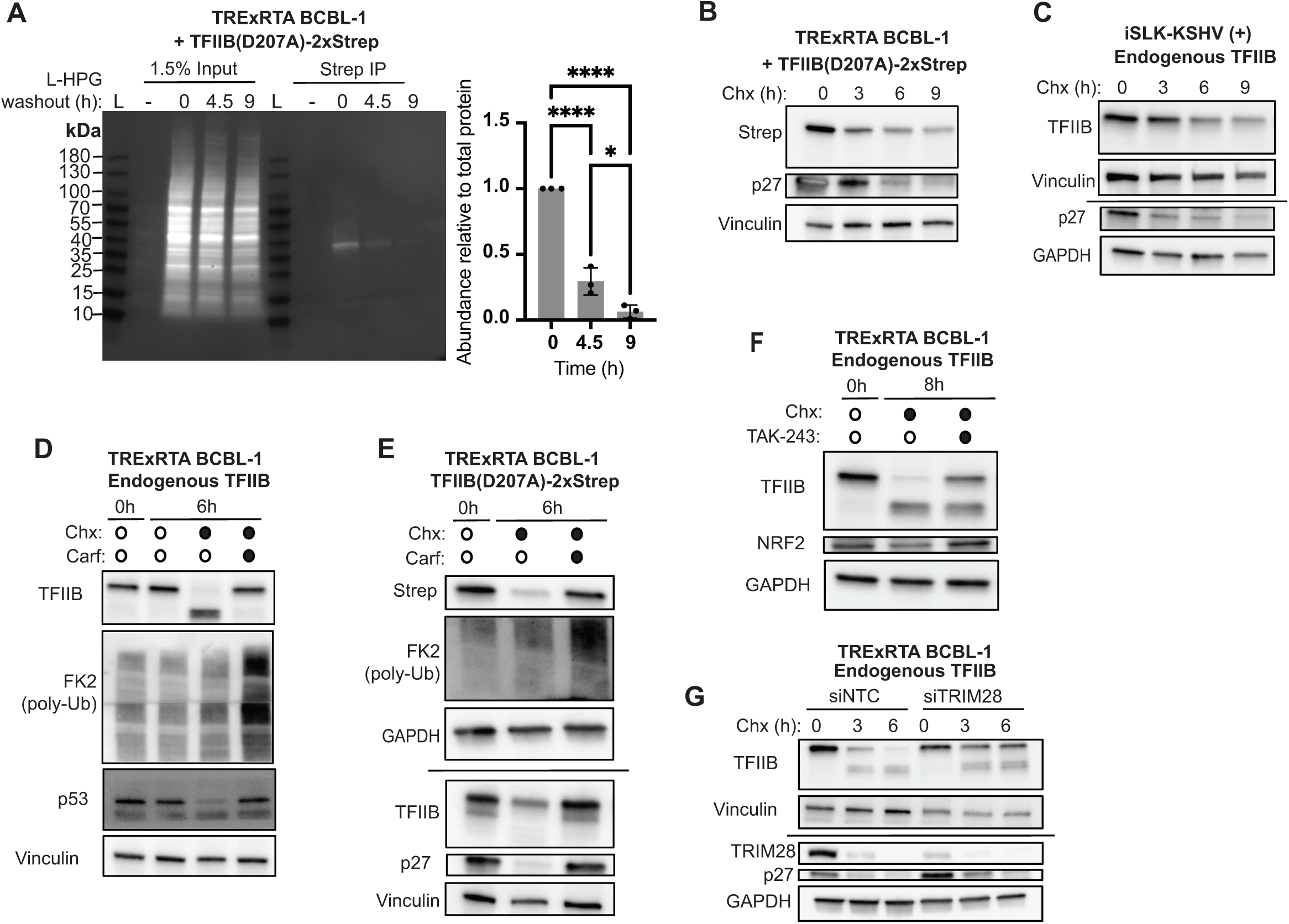
TFIIB is rapidly degraded through the proteasome in both stressed and unstressed cells. **(A)** TRExRTA BCBL-1 cells expressing TFIIB(D207A)-2xStrep protein were pulse-labelled for 18 h with L-HPG, harvested at the indicated timepoints post-chase for strep pulldown, then labelled with picolyl-azide-sulfo-Cy5. For quantification, strep signal was normalized to input signal and the 0 h timepoint and plotted. **** P < 0.0001, * P = 0.0103; one-way ANOVA with Tukey multiple comparison test. **(B)** TRExRTA BCBL-1 cells expressing TFIIB(D207A)-2xStrep were treated with 100 μg/mL cycloheximide (Chx) and harvested at indicated timepoints for western analysis. p27 is a control for Chx activity and vinculin is a loading control. **(C)** iSLK-KSHV(+) BAC16 cells were treated with 250 μg/mL Chx, harvested at the given timepoints and western blotted for the indicated proteins. Vinculin and GAPDH probed with mouse antibody serve as loading controls. **(D)** TRExRTA BCBL-1 cells or **(E)** TRExRTA BCBL-1 cells expressing TFIIB(D207A)-2xStep were treated with 100 μg/mL Chx for 6h with or without 5 μM carfilzomib proteasome inhibitor (carf) and endogenous TFIIB and strep-tagged protein analyzed by western blotting. p53 and p27 are positive controls for Chx activity and FK2 marks poly-ubiquitylated protein as a positive control for proteasomal inhibition; GAPDH probed with mouse antibody is a loading control. **(F)** TRExRTA BCBL-1 cells treated with 100 μg/mL Chx for 8 h with or without 10 μM TAK-243, a UBA1-targetting inhibitor of ubiquitylation, and western blotted with the indicated antibodies. NRF2 is a control for translation and ubiquitylation inhibition, and GAPDH probed with rabbit antibody is a loading control. **(G)** Western blots of TRExRTA BCBL-1 cells treated with 18 h of TRIM28 or nontargeting control (siNTC) siRNAs, followed by 100 μg/mL Chx treatment for the indicated times. Same loading controls as in (*C*).

The other major proteolysis pathway is the ubiquitin proteasome system, and treatment of cells with carfilzomib, a potent and irreversible inhibitor of the 26S proteasome (40), fully restored TFIIB levels in cycloheximide-treated cells (**Fig. 2D**). However, it also inhibited caspase cleavage of TFIIB, making it difficult to determine if TFIIB rescue is due to proteasomal inhibition or indirect inhibition of caspase activity. We therefore tested the caspase-resistant TFIIB(D207A) mutant and found it to be similarly stabilized in cells treated with cycloheximide and carfilzomib (**Fig. 2E)**, establishing proteasomal turnover as an additional protein degradation mechanism for full-length TFIIB. TAK-243, a UBA1-targeting inhibitor of the ubiquitin cascade (41), also increased the levels of full-length TFIIB without impairing caspase cleavage, further supporting a ubiquitin-dependent TFIIB degradation pathway (**Fig. 2F**). A recent substrate-trapping study identified TFIIB as a target of the E3 ligase TRIM28, also known as KAP-1/TIF1β (39). To evaluate whether the TRIM28 E3 ligase reduces the levels of full-length TFIIB in BCBL-1 cells, we used siRNAs to knock down TRIM28. Indeed, compared to control siRNA treated cells, TFIIB protein stability was significantly extended in BCBL-1 cells lacking TRIM28 (**Fig. 2G**). TRIM28 depletion did not impact caspase cleavage, indicating that it operates independently of the caspase-mediated depletion pathway. This effect was specific to TRIM28, as knockdown of TRIM24, a related protein that complexes with TRIM28 (42), did not impact TFIIB levels (***SI Appendix,* Fig. S2D**). Thus, TFIIB abundance is controlled by a combination of TRIM28-mediated proteasomal turnover and caspase cleavage.

### KSHV mitigates TFIIB RNA and protein loss by preventing TFIIB proteasomal degradation

Upon reactivation, KSHV broadly restricts host gene expression by expressing a viral nuclease (SOX) that promotes widespread degradation of cytoplasmic mRNA (14, 15), including TFIIB mRNA (**Fig. 3A**). Thus, KSHV reactivation should in theory lead to similarly robust TFIIB depletion as cycloheximide treatment since the reduced mRNA abundance would restrict new protein synthesis. Yet, lytically reactivated BCBL-1 cells maintain modest levels of TFIIB protein (**Fig. 3B**), suggesting that KSHV has a mechanism to preserve a pool of TFIIB for viral use.

**Figure 3.**
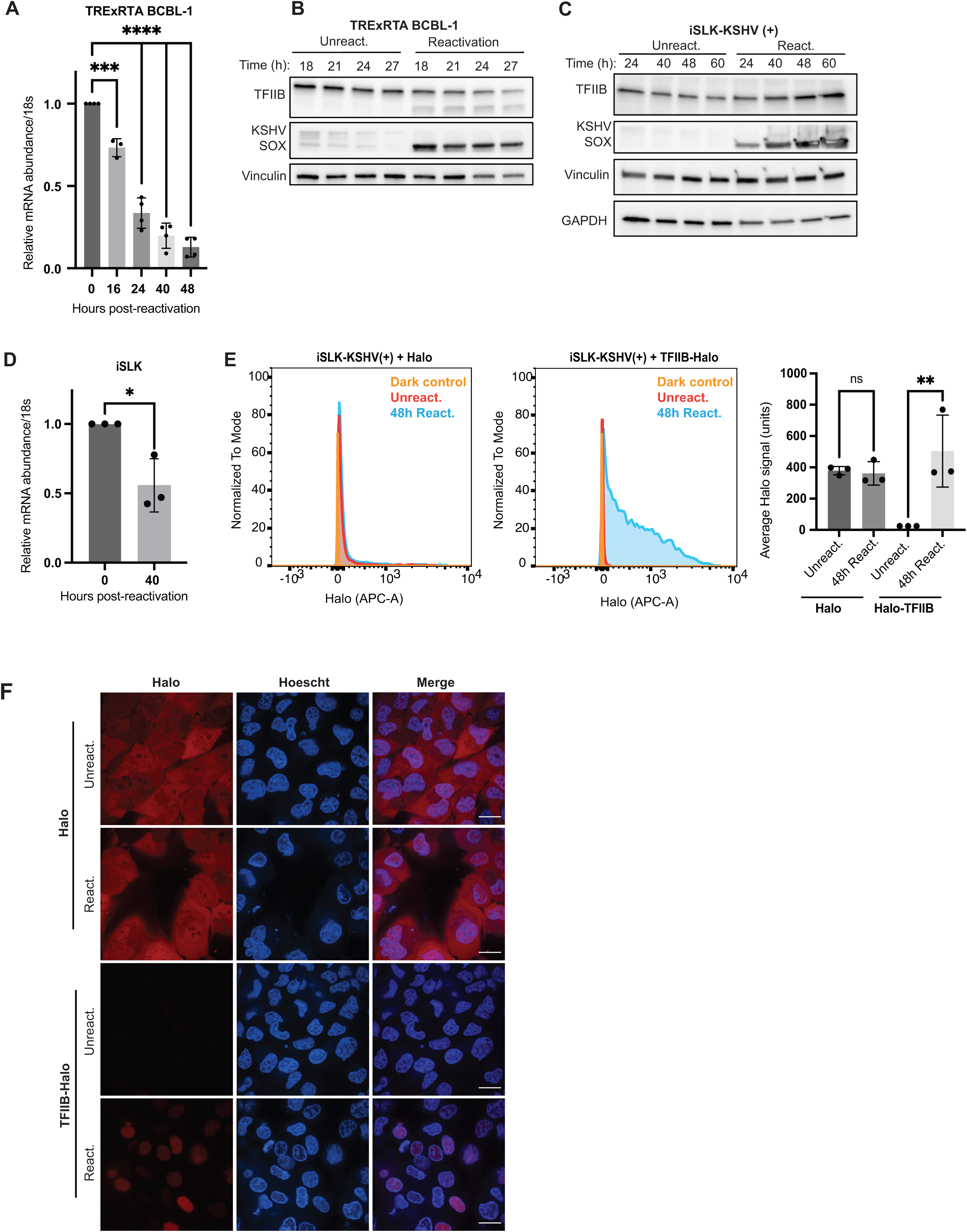
KSHV infection buffers against TFIIB loss. **(A)** RT-qPCR of TFIIB normalized to 18S RNA harvested from reactivated TRExRTA BCBL-1 cells at the given timepoints. **** P < 000.1, *** P = 0.0008; one-way ANOVA with Tukey multiple comparison test. **(B)** Western analysis of endogenous TFIIB in TRExRTA BCBL-1 cells or **(C)** iSLK-KSHV(+) BAC16 cells at timepoints post-reactivation or mock. KSHV SOX is an early gene and vinculin, and GAPDH probed with mouse antibody is a loading control. **(D)** As in (*A*) but in iSLK-KSHV(+) BAC16 cells. * P = 0.01636; Student’s t-test. **(E)** Flow analysis of iSLK-KSHV(+) BAC16 cells transduced with Halo or TFIIB-Halo and mock or 48 h reactivation with doxycycline (dox) and sodium butyrate; Halo was labeled with JF646-Halo dye and measured by signal in the APC-A channel. Fluorescence distributions are shown for a representative replicate and mean APC-A quantification for all samples is shown on the right. ns-nonsignificant, ** P = 0.0057; One-way ANOVA with Tukey multiple comparison test. **(F)** Confocal microscopy of iSLK-KSHV(+) BAC16 cells transduced with Halo protein or TFIIB-Halo, +/- viral reactivation. Hoescht 3342 delineates nuclei. Scale bar = 20 μm.

Indeed, we found that the steady state levels of TFIIB protein increased across a time course of KSHV reactivation in the iSLK cells (**Fig. 3C**), even though TFIIB mRNA abundance decreased (**Fig. 3D)**. These iSLK cells are latently infected with KSHV and contain an integrated, dox-inducible copy of the RTA lytic transactivator that, together with sodium butyrate treatment, promotes viral reactivation into the lytic cycle. To validate this phenotype using an orthogonal assay, we stably transduced iSLK cells with Halo-tagged TFIIB or a Halo only control for visualization by flow cytometry and confocal microscopy. Upon KSHV lytic reactivation, TFIIB-Halo signal increased ∼25-fold, whereas no change was detected for Halo alone (**Fig. 3E)**. The increased TFIIB-Halo signal during reactivation was localized to the nucleus, while expression of the Halo control protein was unchanged and localized diffusely throughout the cell (**Fig. 3F**). Both endogenous TFIIB and Halo-TFIIB levels remained largely unchanged upon dox and sodium butyrate treatment of control uninfected iSLK cells (that still contained dox-inducible RTA) (***SI Appendix*, Fig. S3A-B**). Thus, lytic reactivation is the main driver of increased TFIIB abundance.

We next compared the half-life of TFIIB protein in unreactivated versus lytically reactivated cells upon inhibiting new protein synthesis with cycloheximide. The stability of full-length TFIIB was markedly increased upon lytic viral reactivation in both iSLK and BCBL-1 cells (**Fig. 4A-B**). This was specifically induced by KSHV reactivation, as etoposide treatment did not similarly extend TFIIB’s half-life (***SI Appendix*, S4A).** The increase in TFIIB levels upon reactivation was maintained in cells treated with the KSHV DNA replication inhibitor phosphonoacetic acid (PAA), which also blocks all subsequent lytic cycle events **(Fig. 4C; *SI Appendix*, Fig. S5A**). Thus, TFIIB stabilization occurs during the early phase of the lytic cascade. TFIIB stabilization was not a consequence of impaired caspase cleavage, as we also observed stabilization of the caspase-resistant TFIIB(D207A) mutant (**Fig. 4D**). This instead suggests that reactivation leads to a defect in its proteasomal decay. While we favor the hypothesis that this stabilization involves counteracting TRIM28, we have observed neither differences in TRIM28 abundance in reactivated cells nor a role for specific KSHV-induced TRIM28 phospho-regulation in the TFIIB stabilization phenotype (***SI Appendix*, Fig. S4B-F**). Thus, more complex regulation with other modifications and/or E3 ligases may be involved and will be the focus of future work.

**Figure 4.**
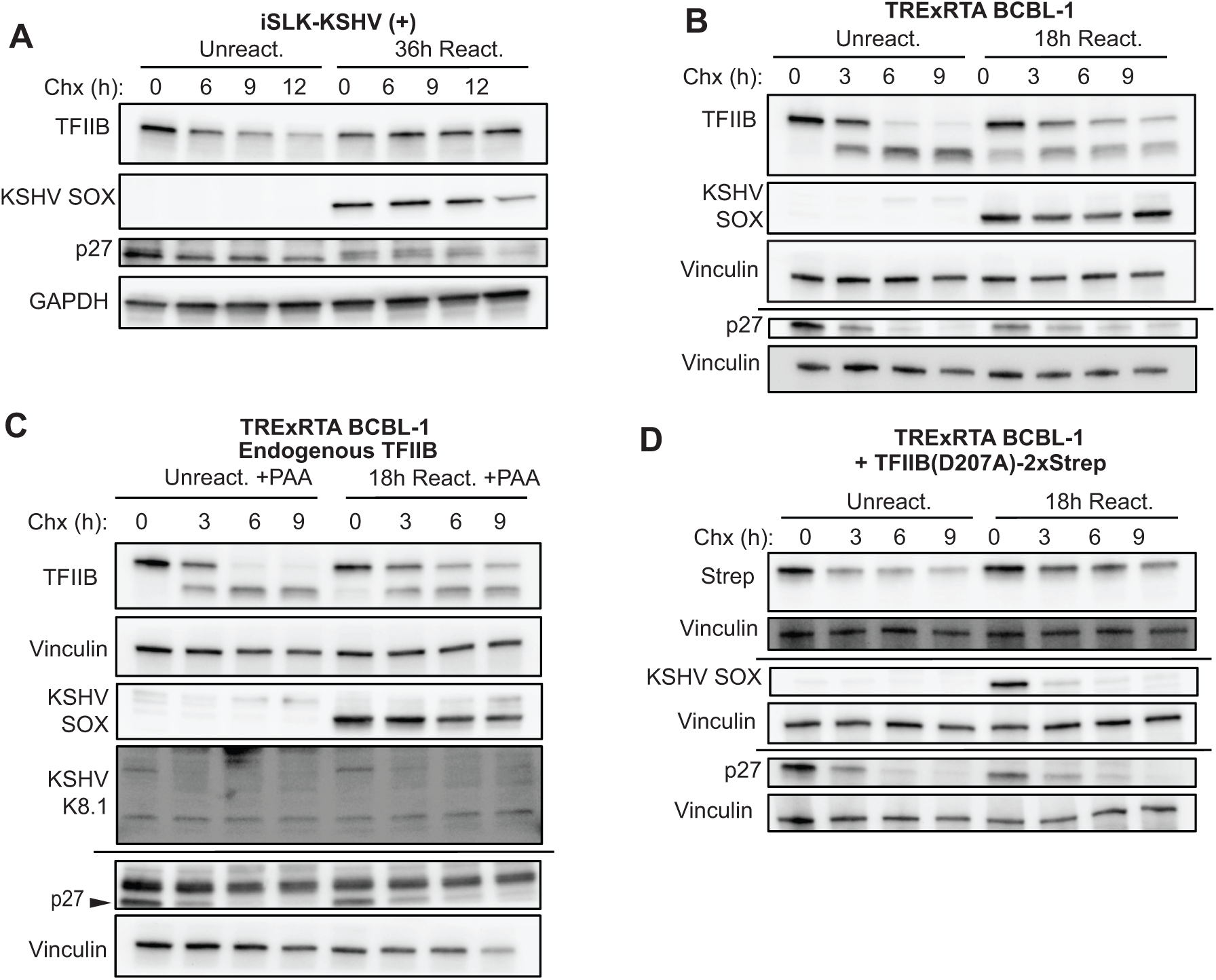
KSHV infection impairs TFIIB proteasomal degradation early in infection. **(A)** iSLK-KSHV(+) BAC16 cells +/- reactivation for 36 h, treated with 250 μg/mL cycloheximide (Chx) for the indicated timepoints, then western blotted with the indicated antibodies. KSHV SOX is a control for infection, p27 is a control for translation inhibition, and GAPDH probed with rabbit antibody is a loading control. **(B)** TRExRTA BCBL-1 cells were mock-treated or reactivated for 18 h, treated with 100 μg/mL Chx, and western blotted for the indicated proteins. Vinculin is a loading control. **(C)** Western blots of TRExRTA BCBL-1 cells +/- 18 h reactivation and treatment with 1mM of the viral DNA replication inhibitor phosphonoacetic acid (PAA) followed by a timecourse with 100 μg/mL Chx. Early viral gene SOX expression is not affected by inhibition of DNA replication, but late gene K8.1 is dependent on DNA replication and thus not expressed. Black arrow indicates appropriate p27 band under non-specific antibody signal. GAPDH probed with mouse antibody is a loading control. TRExRTA BCBL-1 cells expressing TFIIB(D207A)-2xStrep were mock-treated or reactivated for 18 h, treated with 100 μg/mL Chx, and western blotted for the indicated proteins. Vinculin is a loading control. **(D)** As in (*B*) but with TRExRTA BCBL-1 cells expressing TFIIB(D207A)- 2xStrep.

### KSHV gene expression is impaired upon TFIIB depletion

Finally, since KSHV requires RNAPII activity for viral transcription, we asked whether retention of a pool of TFIIB was important for KSHV gene expression. We reduced the TFIIB levels during KSHV lytic reactivation using TFIIB-targeting siRNAs in BCBL-1 cells (**Fig. 5A; *SI Appendix*, Fig. S5A-B**). RNA-seq revealed decreased expression of nearly all KSHV genes compared to a control siRNA at 48 h post-lytic reactivation (**Fig. 5B-C**; ***SI Appendix* Fig. S5C-D**). This was not simply due to decreased viral reactivation, since even at 24 h post-reactivation in TFIIB knockdown conditions KSHV gene expression appeared mis-regulated, with reduced transcript levels of genes normally expressed early and increased levels of some genes usually expressed only late in infection (***SI Appendix,* Fig. S5D**). As anticipated, the majority of differentially expressed cellular genes were also decreased upon TFIIB knockdown compared to control siRNAs, which we were able to evaluate as samples were normalized to spike-in controls (**Fig. 5B; *SI Appendix*, Fig. S5C**). Conversely, expression of the majority of the KSHV genes was modestly increased in RNA-seq data from cells stably expressing the cleavage-resistant TFIIB(D207A)-2xStrep versus those expressing cleavable, wild-type TFIIB-2xStrep (**Fig. 5D**; ***SI Appendix*, Fig. S5E)**. Though no individual gene varied significantly, the ensemble of relative changes in the viral transcriptome was significantly upregulated in TFIIB(D207A)-2xStrep expressing cells over TFIIB-2xStrep (**Fig. 5E**). TFIIB knockdown also led to lower levels of viral DNA replication, as measured by viral DNA qPCR at 48 hours post-reactivation (**Fig. 5F**). Thus, KSHV requires sufficient TFIIB levels for proper viral gene expression and has evolved a mechanism to preserve a pool of TFIIB under conditions where it would normally be depleted.

**Figure 5.**
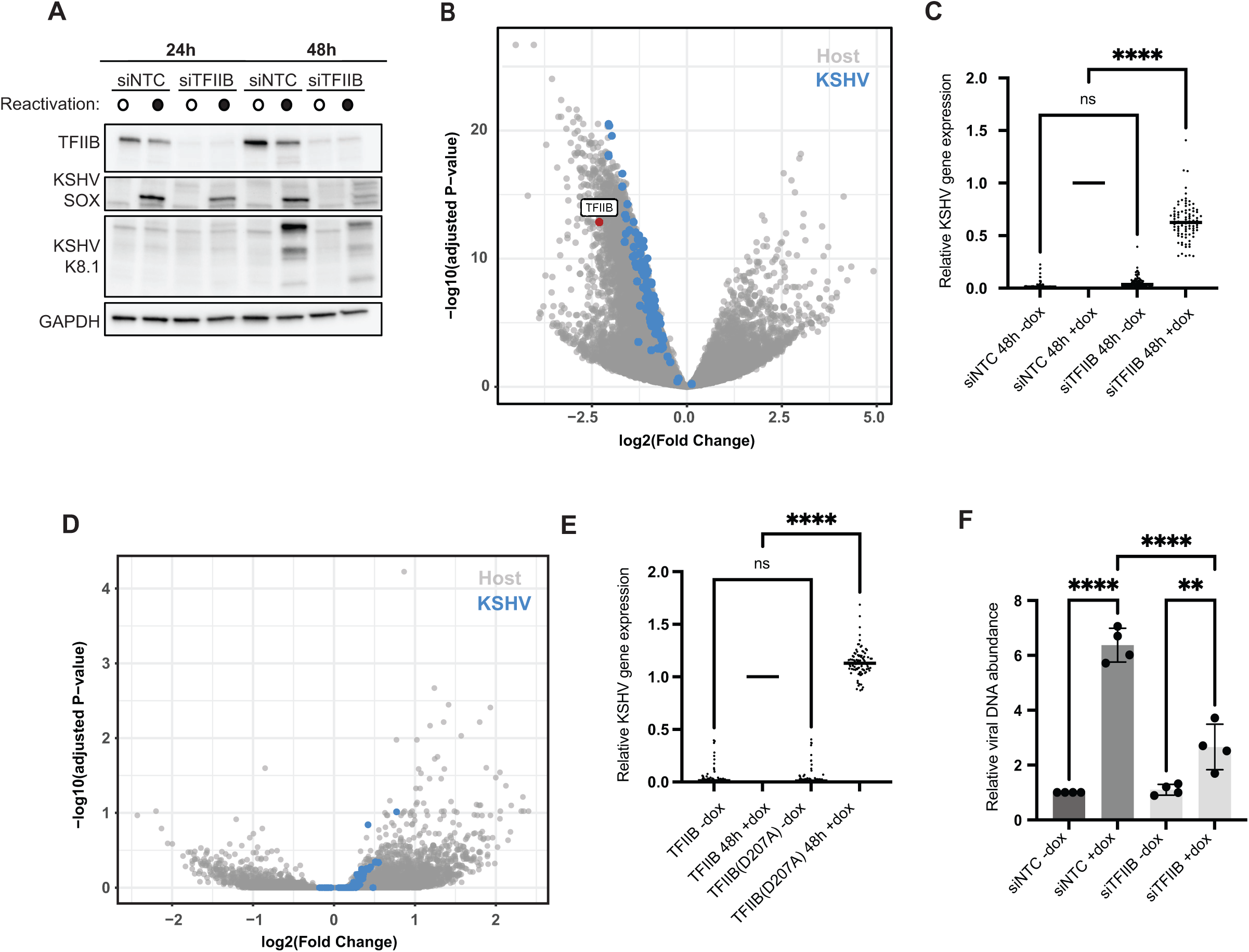
KSHV requires TFIIB for optimal gene expression. **(A)** Western blotting to confirm TFIIB knockdown in TRExRTA BCBL-1 cells nucleofected with TFIIB siRNA (siTFIIB) or non-targeting siRNA (siNTC) followed by 24 h or 48 h of doxycycline (dox)-induced reactivation or mock. SOX is a viral early gene and K8.1 is a viral late gene. GAPDH probed with mouse antibody is a loading control. **(B)** Volcano plot of differentially expressed genes from the RNA-seq analysis of TRExRTA BCBL-1 treated with either TFIIB siRNA (siTFIIB) or nontargeting siRNA (siNTC) and reactivated for 48 h. Cellular genes are in gray and KSHV genes are highlighted in blue. **(C)** Relative gene expression for KSHV genes in TRExRTA BCBL-1 cells treated with siNTC or siTFIIB knockdown +/- doxycycline (dox)-induced reactivation. Calculations were derived from viral RNA-seq gene counts normalized to siNTC with 48 h reactivation (siNTC 48 h reactivation was normalized to 1). ns- nonsignificant, **** P < 0.0001; one-way ANOVA with Tukey multiple comparison test. **(D)** Volcano plot of differentially expressed genes from the RNA-seq analysis of TRExRTA BCBL-1 expressing either TFIIB-2xStrep or TFIIB(D207A)-2xStrep and reactivated for 48 h. Cellular genes are in gray and KSHV genes are highlighted in blue. **(E)** Relative gene expression for KSHV genes in TRExRTA BCBL-1 cells expressing either TFIIB-2xStrep or TFIIB(D207A)- 2xStrep +/- dox-induced reactivation. Calculations were derived from viral RNA-seq gene counts normalized to TFIIB-2xStrep with 48 h reactivation (which itself was normalized to 1). ns-nonsignificant, **** P < 0.0001; one-way ANOVA with Tukey multiple comparison test. **(F)** Relative viral DNA abundance was measured by qPCR in siNTC or siTFIIB treated TRExRTA BCBL-1 cells with 48 h post dox-induced reactivation as measured by viral late gene K8.1 promoter amplification normalized to cellular TLDC1 promoter amplification by qPCR. **** P < 0.0001, ** P = 0.0060; one-way ANOVA with Tukey multiple comparison test.

## Discussion

Here we uncovered a two-pronged mechanism by which cells control abundance of the RNAPII general transcription factor TFIIB. In unstressed cells, TFIIB is continually turned over through ubiquitin-mediated proteolysis. A labile core GTF may facilitate rapid transcriptional tuning as cells encounter environmental perturbations. In response to cell stress, TFIIB is further depleted by caspase activity, which we hypothesize leads to preferential suppression of the most TFIIB-dependent genes. In apoptotic cells, these include genes involved in basal pro-growth and pro-survival pathways. This may also be a strategy by which cells can limit DNA viruses’ access to gene expression machinery. However, KSHV counteracts this response by inhibiting TFIIB proteasomal degradation, enabling robust viral gene expression.

### Stress reshapes transcription by targeting RNAPII protein machinery

Rapid responses to environmental stimuli are essential for cell viability. Canonically, this involves dramatic alterations to transcription and translation. In the cytoplasm, there is a general arrest of translation initiation and sequestration of mRNA into stress granules (reviewed by Advani and Ivanov(43)). Similarly, in the nucleus, repression of bulk transcription may play a key role in shifting resources to hyper-activate stress response or antiviral genes, as well as potentiate cellular shutdown when cell death is triggered. Several mechanisms exist to post-translationally target and control the RNAPII holoenzyme, including ubiquitin-mediated degradation of the RNAPII catalytic subunit (RPB1/POLR2A) during genotoxic stress (4, 44, 45). This promotes temporary transcriptional shutdown of short genes, which are often pro-growth factors and oncogenes (4).

Analogously, we found that TRAIL-treated cells expressing a caspase-resistant version of TFIIB upregulated several factors that favor cell survival, including the anti-apoptotic BCL2 regulator. Although the number of induced genes was relatively small, we nonetheless found this shifted a proportion of cells from apoptotic cell death to necrotic death. Perhaps the inability to shut down expression of these TFIIB-dependent genes prevents proper execution of apoptosis and results in an inappropriate, more inflammatory (46) form of cell death. It is thus possible that the abundance of TFIIB and other RNAPII machinery in a cell are balanced to determine cell fate. In keeping with this idea, TFIIB-dependent metabolic genes are commonly dysregulated in cancer cells (47), and rapid TFIIB proteasomal turnover also occurs during cellular differentiation of F9 cells, along with depletion of other RNAPII GTFs (48). We hypothesize that an appropriate cellular stress response requires a tradeoff from normal TFIIB-dependent transcription to favor expression of stress-responsive pathways.

### TFIIB plays key roles during viral infection

Control of TFIIB, and more broadly RNAPII, is an important component of virus-host interactions (8, 49). RNA viruses that do not use cellular RNAPII for gene transcription can selectively cripple host gene expression by targeting core GTFs like TFIIB, thereby limiting antiviral gene expression (50, 51). For example, during Thogoto virus infection, the viral ML protein dramatically relocalizes TFIIB to the cytoplasm as an immune evasion strategy; this impairs both the innate immune response and apoptotic pathways in HeLa cells (50), suggesting a context-dependent regulatory role for TFIIB abundance. On the other hand, nuclear replicating DNA viruses must hijack cellular transcriptional machinery for viral gene expression. Downregulation of TFIIB here may be an antiviral response geared towards restricting viral access to functional transcriptional machinery while maintaining sufficient levels to activate anti-viral genes. To counter, DNA viruses must evolve mechanisms to preserve GTF access for viral needs. Many DNA viruses encode proteins that interact with and recruit TFIIB to viral promoters. Examples from other *Herpesviridae* include herpes simplex virus 1 VP16 (52, 53), human cytomegalovirus IE2 (54), and Epstein-Barr virus EBNA2 (55). These interactions are presumably important for viral fitness as knockdown of TFIIB significantly impairs HSV-1 replication (56), similar to our observations with KSHV. Whether these and other DNA viruses also block proteasome-mediated turnover of TFIIB will be interesting to explore in the future.

A key open question is how KSHV coordinates TFIIB stabilization. While it is possible that an early viral protein directly interacts with and stabilizes TFIIB during KSHV reactivation, a previously published KSHV-human protein-protein interaction map failed to detect any such interaction (57), suggesting the mechanism of stabilization is indirect. Given that TRIM28 has E3 ligase activity that targets TFIIB for turnover (39), we favor a model in which one or more KSHV genes antagonizes TRIM28 activity. TRIM28 is extensively modified by mammalian viruses, with many examples from the *Herpesviridae* (58), where it has roles in both viral latency and the switch to lytic replication (59–64). Protein abundance and phosphoregulation of TRIM28 do not appear to underly enhanced TFIIB stability during KSHV reactivation, but a more extensive survey of TRIM28 modifications during the lytic viral cycle may yield other specific modifications to examine. It is possible that additional E3 ligases also regulate TFIIB and may be blocked during the KSHV lytic cycle. Future studies addressing these, and other possibilities should further our understanding of TFIIB and RNAPII holoenzyme regulation, which is central to how cells elicit broad transcriptional responses when under threat.

## Methods

A comprehensive list of materials and identifiers may be found in ***SI Appendix;* Table S2**

### Plasmids and primers

Primers and relevant DNA sequences are listed in ***SI Appendix;* Table S3**.

### Cloning

TFIIB constructs were cloned from TRExRTA BCBL-1 cell cDNA. The DNA sequence was modified to preserve the amino acid sequence but render constructs resistant to Dharmacon OnTarget and Accell human GTF2B siRNAs; this sequence was synthesized for cloning by Twist Bioscience and used for all subsequent plasmid generation. For strep-tagged constructs, TFIIB was subcloned into XhoI site of pcDNA4/TO-2xStrep (pCDNA4/TO Invitrogen vector modified to include a C-terminal 2xStrep tag) using InFusion cloning (Takara Bio). TFIIB-2xStrep was then amplified and subcloned into the Sfi1 site of a sleeping beauty plasmid pSBbi-GP (Addgene #60511, gift from Eric Kowarz) derivative lacking EGFP (gift from Nicholas Ingolia, UC Berkeley) by InFusion cloning to make the TFIIB-2xStrep expression vector. Derivative point mutant plasmids D207A, S65A, and S65E were generated by inverse PCR (described in Silva et al.(65)). Sleeping beauty vectors were used with sleeping beauty transposase plasmid (gift from Nicholas Ingolia, UC Berkeley); the sleeping beauty system was described in Kowarz et al.(66).

Lentivectors driving HaloTag or TFIIB-Halo expression were generated by amplifying TFIIB and HaloTag for C-terminal insertion by InFusion cloning into EcoRV-digested pLJM1 (Addgene plasmid #19319, gift of David Sabatini, modified to remove EGFP and replace puromycin resistance with a blasticidin resistance gene from pSBtet-Bla, Addgene plasmid #60510, gift of Eric Kowarz). TFIIB was immediately followed by a short, flexible GGSGGGS linker before the HaloTag protein. Halo only vector was made by cloning out TFIIB with inverse PCR and T4 PNK/T4 ligase religation. Lentivectors were co-transfected with packaging plasmids pMD2.G (Addgene plasmid #12259) and psPAX2 (Addgene plasmid #12260), gifts of Didier Trono.

All plasmids constructed in this study and the HA-Kaposin B and HA-EGFP vector were validated by full-plasmid sequencing and have been deposited on Addgene (accession numbers pending).

### Cell line generation, maintenance, and reactivation

All cell lines were maintained at 37°C and 5% CO_2_ in a humidity-controlled incubator. TRExRTA BCBL-1 cells (generously provided by the lab of Jae Jung) contain a doxycycline-inducible copy of the KSHV lytic transactivator RTA.(26) Cells were maintained in RPMI 1640 media (Gibco) supplemented with 10% Tet-free fetal bovine serum (FBS; via UC Berkeley Cell Culture Facility), 1X GlutaMAX (Gibco), and 10 μg/mL hygromycin B (Invitrogen). TRExRTA BCBL-1 derivative cell lines were also passaged with 1 μg/mL puromycin (Gibco) for selection. Cells were kept at densities between 0.5-2x10^6^ cells/mL in non-tissue culture treated dishes. Unless otherwise noted, cell concentration was ∼1x10^6^ cell/mL when reactivated with 1 μg/mL doxycycline or administered other drug treatment. TRExRTA BCBL-1 cells line derivatives were generated by nucleofection with the Neon Transfection System (Invitrogen) according to the manufacturer’s instructions with the following parameters: 5x10^6^ low-passage cells were pulsed (1350V, 1x40ms pulse-length) with 2 μg DNA in a 100 uL tip. Sleeping beauty plasmids containing TFIIB-2xStrep and point mutant derivatives were delivered with a 1:20 ratio of transposase to delivery vector. Cells were recovered for 24 h prior to commencing 1 μg/mL puromycin selection.

iSLK-BAC16 cells(67–69) are renal carcinoma cells that contain the KSHV genome on bacterial artificial chromosome (BAC) and a doxycycline-inducible copy of RTA. Control iSLK cells similarly contain dox-inducible RTA but lack the KSHV BAC. Cells were maintained in DMEM media supplemented with 10% FBS (Peak Serum), 1 mg/mL hygromycin B, and 1 μg/mL puromycin (iSLK cells without BAC16 were propagated identically but without hygromycin). iSLK-BAC16 derivatives were also maintained in 10 ug/mL blasticidin (Gibco). For lytic reactivation, cells were plated at ∼0.25-0.5 x10^6^ cells per well the day before treatment and reactivated at 70-80% confluency with 5 μg/mL doxycycline (Millipore Sigma) and 1 mM sodium butyrate (Sigma). iSLK and iSLK-BAC16 cell line derivatives were generated using 2^nd^ generation lentivirus as follows: 1x10^6^ Lenti-X 293T cells (Clonetech/Takara via the UC Berkeley Tissue Culture Facility; maintained in DMEM media (GIBCO) with 10% FBS) were transfected using *Trans*IT-X2 or *Trans*IT-LT1 Transfection Reagent (Mirus; using the manufacturer’s protocol) with 250 ng pCMV-VSG, 1250 ng psPAX2, and 1250 ng lentivector. Lentivirus was harvested at 48 h post-transfection with a 0.45 μM filter and frozen at -80°C in a total volume of 5 mL of media. iSLK-BAC16 cells at ∼80% confluency were incubated with 2 mL regular media and 1 mL of thawed lentivirus for 24 h, at which point the media was refreshed. Cells recovered for 24 h prior to selection with 10 μg/mL blasticidin.

### siRNA knockdowns and nucleofections

3x10^6^ TRExRTA BCBL-1 cells in 100 μL of resuspension buffer T (Invitrogen) were mixed with 3 μL of 100 μM siRNA stock of the following Dharmacon siRNAs: On-Target Plus Non-targeting control pool and Accell Non-targeting control pool for non-targeting control cells, or On-Target Plus human GTF2B siRNA SmartPool and Accell human GTF2B siRNA SmartPool for TFIIB knockdown cells. Cells were nucleofected with the Neon Transfection System (Invitrogen) according to the manufacturer’s instructions at 1350V, 1x40ms pulse-length. Two rounds of siRNA delivery were performed on each pool of cells, 24 h apart. Further experimentation (e.g., reactivation) took place 24 h after the last siRNA delivery.

Caspase-3 knockdown was performed as described for TFIIB using the following Dharmacon siRNAs: Accell Human CASP3 siRNA SmartPool and ON-Target plus Human CASP3 siRNA SmartPool, with appropriate control pools. This experiment was modified with only one siRNA treatment administered 48 h prior to cycloheximide treatment and further experimentation.

TRIM28 and TRIM24 knockdowns were performed similarly with the following modifications: 3x10^6^ TRExRTA BCBL-1 cells in 100 μL of resuspension buffer T (Invitrogen) were nucleofected once with 2 μL of 100 μM of either Dharmacon human TRIM28 Accell SmartPool siRNA or OnTarget Plus human TRIM24 SmartPool. Cycloheximide treatment and harvesting took place 18 h after nucleofection.

HA-Kaposin B and HA-EGFP expression experiments were performed using the same nucleofection parameters for one-time delivery of 3 μg HA-Kaposin B or 1.5 μg HA-EGFP vectors (to better match protein expression levels) into 3x10^6^ TRExRTA BCBL-1 cells for each condition 16 h prior to further experimentation.

### Western blotting

Cells were collected and washed by either centrifugation at 300-1,000 x g for 3 min (suspension cells) or scraping in 1 mL of PBS. Cells were re-pelleted with a high spin for 20s in a microfuge tube and lysed in 1% SDS RIPA buffer (1% SDS, 1% NP40, 150 mM NaCl, 50 mM Tris-HCl pH 8, 0.5% sodium deoxycholate by mass) with Halt Protease and Phosphatase Inhibitor Cocktail (Thermo Scientific). Cell lysates were stored at -80°C until further processing. Lysates were sonicated (qSonica Ultrasonicator with cuphorn for 1 min at 100A, 3 s on/17 s off) and treated with 1% Benzonase nuclease (Millipore). Lysates were clarified by centrifugation at 21,000 x g for 10 minutes at 4°C and quantified by Pierce 660 assay according to the manufacturer’s cuvette protocol, using 0.8% Triton-X (MilliQ H_2_O was used to dilute lysates). 20-30 μg whole cell lysate was denatured in Laemmli Buffer (Bio-Rad), resolved by SDS-Page, and transferred to PVDF membrane (Bio-Rad) with a Trans-Blot Turbo (Bio-Rad, Mixed Molecular Weight protocol). Western blotting was performed with Tris-buffered saline and 0.2% Tween-20 (TBS-T) using PageRuler Prestained protein ladder (10-180 kDa formulation; ThermoFisher) and the following primary and secondary antibodies: TFIIB (clone 2F6A3H4, Cell Signaling-4169, 1:1000), GAPDH (anti-mouse, clone 6C5, Thermofisher-AM4300, 1:5000 or anti-rabbit, clone 14C10, Cell Signaling-2118, 1:1000), p27 (ProteinTech-15086-1-AP, 1:1000), TRIM28 (Cell Signaling-4123, 1:1000), TRIM28-P(S824) (Cell Signaling-4127, 1:1000), TRIM28-P(S473) (clone 11G10SC, Biolegend-654101, 1:500-1:1000), TRIM24 (clone E93TN, Cell Signaling-79030, 1:1000), Vinculin (Abcam-ab91459, 1:1000), FK2 (Enzo-BML-PW8810-0100, 1:1000), HA (clone C29F4, Cell Signaling-3724, 1:1000), NRF2 (ProteinTech-16396-1-AP, 1:000), Caspase-3 (Cell Signaling-9662, 1:1000), Caspase-8 (clone E7, Abcam-ab32397, 1:1000), γH2AX (Bethyl-A300-081A-M, 1:1000), p53 (clone 1C12, Cell Signaling-2524, 1:1000), Goat Anti-Mouse IgG(H+L) Human ads-HRP (Southern Biotech, 1:5,000), Goat Anti-Rabbit IgG(H+L) Human ads-HRP (Southern Biotech, 1:5000), Strep-Tag II Antibody HRP Conjugate (Sigma Aldrich-71591-3, 1:2000). Rabbit anti-K8.1 (gift of Denise Whitby, 1:10000)(70) and Rabbit anti-SOX (Yenzyme, YZ6540 (cycle 1), 1:5000)(71) were described previously. Blots were developed with either Clarity Western ECL Substrate (Bio-Rad) or Radiance Plus Femtogram-sensitivity HRP Substrate (Azure Biosystems). Blots were imaged on a Chemidoc MP imager (Bio-Rad) and prepared for publication using Bio-Rad Image Lab (version 6.0.1) using the autoscale feature to maintain consistency unless otherwise noted. Blots shown are representative of a minimum of 3 biological replicates at the same conditions and/or timepoints.

### qPCR

#### RNA

Cells were harvested as described for westerns and total RNA was collected in 1 mL TRIzol (Invitrogen) and isolated by 1 x 200 uL chloroform extraction as outlined in Toni et al. (72), including precipitation in 500 uL isopropanol with 1% glycogen and 10% 3 M sodium acetate. Purified RNA was treated with Turbo DNase (Thermofisher) and reverse-transcribed with AMV-RT (Promega) and random 9-mer primers. cDNA was quantified by qPCR with a QuantStudio3 Real-Time PCR machine and transcript-specific primers and iTaq UniverSYBR Green Supermix (Bio-Rad). RNA was normalized to 18S rRNA levels and corrected for primer efficiency.

#### vDNA quantitation

DNA was isolated from cells using QiaAMP Blood Mini kit (Qiagen) according to the manufacturer’s protocol. Equivalent concentrations of total DNA were loaded for quantitation by QuantStudio3 Real-Time PCR (Applied Biosystems) with iTaq UniverSYBR Green Supermix (Bio-Rad). Fold viral DNA replication changes were assessed using primers specific to the KSHV K8.1 promoter normalized to the human TLDC1 promoter. A dilution series of total DNA verified that there was no inhibition of the PCR reaction at the concentrations tested.

### L-Homoproparglyglycine and click labeling

TREx-RTA BCBL-1 cells were washed with DPBS (Gibco) and 10x10^6^ cells per condition were seeded at 1 x10^6^ cells/mL in normal RPMI media with or without methionine. Cells without methionine were pulsed with 50 μM Click-It L-Homopropargylglycine (L-HPG) (Thermofisher) for 18h, which is a methionine analog with an alkyne for labeling via click chemistry. Next, cells were pelleted, washed once with DPBS, and chased with 10 mL of fresh methionine-containing RPMI. A 0 h sample and cells with continuous methionine were harvested at this point. Additional samples were harvested at the indicated post-chase timepoints. Cells were lysed in 1% SDS RIPA buffer (1% NP40, 1% SDS, 150 mM NaCl, 50 mM Tris-HCl pH 8, 0.5% sodium deoxycholate by mass) with 1x Halt Protease and Phosphatase inhibitor cocktail (Thermo Scientific) and rotated at 4°C for 1 h, followed by treatment with Benzonase Nuclease (Millipore). Protein concentration was quantified with Pierce 660 as described for Western blots. 1 mg of protein was diluted in Immunoprecipitation (IP) lysis buffer to 1 mL (50 mM Tris pH7.4, 150 mM NaCl, 1mM EDTA, 0.5% NP40, 1x Halt Protease and Phosphatase inhibitor cocktail (Thermo Scientific)) and 1.5% inputs were removed, mixed with Laemmli Buffer (Bio-Rad), and stored at -20°C. Strep protein was immunoprecipitated with 20 uL MagStrep “type3” XT beads (IBA Lifesciences) pre-washed with IP wash buffer (50 mM Tris-HCl pH 7.4, 150 mM NaCl, 1mM EDTA, 0.05% NP40) and rotated overnight at 4°C. After washing beads with additional IP wash buffer, protein was eluted by boiling beads in Laemmli buffer (Bio-Rad) at 95°C (10 min) and resolved by SDS-PAGE as described for Western blots. The click reaction was performed on the PVDF membrane as described by Ohata et al.(73) Briefly, the blot was rinsed twice with methanol, twice with MilliQ H_2_O, and then rocked in the dark, overnight at 4°C in click reagents (1:1 DMSO/H_2_O with 20 μM picolyl-azide-sulfo-Cy5 (Jena Bioscience), 300 μM CuSO_4_, 60 μM tris-hydroxypropyltriazolylmethylamine (THPTA), and 3.0 mM sodium ascorbate). The blot was then washed with methanol, three 5 min DMSO rinses, methanol, and three 70% ethanol rinses before imaging with a ChemiDoc MP (Bio-Rad). Data was quantified by dividing the strep band click signal to overall IP click signal and prepared for publication using Bio-Rad Image Lab (version 6.0.1). Data graphing and analysis were performed in GraphPad Prism 10.10 (264).

### Fluorescence Microscopy

iSLK-BAC16 cell derivatives were plated on glass-bottom tissue culture-treated dishes and reactivated as described earlier. Cells were labelled at least 30 minutes prior to imaging with Halo ligand Janelia Fluor 646 (Promega) at a 1:1,000 dilution of 20 μM stock in DMSO and 10 μM Hoesct dye. 48 h post-reactivation or mock, cells were imaged live using an inverted Nikon Eclipse Ti spinning disk confocal microscope driven by Nikon Imaging Systems (NIS) Elements (v. AR 5.21.03) with a 60× oil immersion objective (Nikon Plan Apo VC 60× Oil DIC N2, 1.4 NA). Single representative images were selected from Z-stacks and image analysis was performed in FIJI(74) (ImageJ v. 2.0.0-rc-69/1.52p), including image merging.

### RNA sequencing and data analysis

For all sequencing experiments, samples were collected as indicated previously and RNA extracted with TRIzol as described for RT-qPCR and quantified by Qubit 3 Fluorometer with High Sensitivity Kit (Invitrogen). Prior to library prep, ERCC RNA Spike-in mix (Invitrogen) was added to all input samples per the manufacturer’s instructions (added for downstream normalization; important for RNA samples with changes in bulk mRNA abundance, as in viral infection.(9) TFIIB knockdown libraries were prepped with KAPA Stranded RNA-seq Kit with RiboErase (HMR) (Roche) using 500 ng of input RNA and amplified for 9 cycles, and TFIIB-2xStrep/TFIIB(D207A)-2xStrep libraries were prepped with KAPA RNA Hyperprep Kit with RiboErase (Roche) using 1,000 ng input RNA and amplified for 7 cycles. Both were fragmented at 94°C for 6 min and ligated with KAPA Unique Dual-Indexed adaptors (Roche). Library quality was checked on an AATI (now Agilent) Fragment Analyzer. Library molarity was measured via quantitative PCR with the KAPA Library Quantification Kit (Roche KK4824) on a BioRad CFX Connect thermal cycler. Libraries were then pooled by molarity and sequenced, targeting at least 25M reads per sample. TFIIB knockdown sequencing was performed on one lane of an Illumina NovaSeq 6000 S1 flowcell with 150 bp paired-end reads. TFIIB-2xStrep/TFIIB(D207A)-2xStrep libraries were sequenced across two lanes of an Illumina NovaSeq X 10B with 150 bp paired-end reads. Fastq files were generated and demultiplexed using Illumina bcl_convert and default settings, on a server running CentOS Linux 7.

Analysis was modified from UC Davis Bioinformatics Core RNA-seq workflow. Sequencing quality was assessed with MultiQC and reads were preprocessed with HTStream version 1.3.0, including deduplication. Genome indices were prepared using STAR (v. 2.7.1a) (75). The human GRCh38.p13 genome assembly (ensembl.org) was indexed with Gencode v43 annotations. Due to overlapping transcripts on the KSHV-BAC16 genome, individual exon coordinates were assigned to the corresponding parent transcript for the virus in a custom gene annotation. Preprocessed reads were aligned with STAR and counts files were generated for transcripts. Raw data and counts tables will be available on the GEO database. For direct analysis of only viral transcripts, viral counts were imported to Microsoft Excel and normalized to the sum of ERCC spike-in counts. Replicates were then normalized to their corresponding mock condition and the replicates averaged. Any transcript with no reads in some or all replicates and conditions was eliminated from further analysis. Reads from *E.coli* genes on the BAC were also removed. Heatmaps were generated using the matplotlib, pandas, and seaborn packages in Spyder (v. 5.3.3). Whole genome and differential gene expression (DGE) analysis was conducted in RStudio (v. 2023.06.2+561) with the Bioconductor package using edgeR with limma/voom developed and described by Law et al.(76). Samples were normalized to ERCC Spike-In and compensated for multiple testing using the Benjamin-Hochberg correction. Genes were annotated with the BioMart Ensembl Genes 109 database (Ensembl). Volcano plots were generated using ggplot2.

### Halo fluorescence measurements

Cells were reactivated or mock treated for 48 h and then labelled at least 30 minutes prior to harvesting with Halo ligand Janelia Fluor 646 (Promega) at a 1:1,000 dilution of 20 μM stock in DMSO. Cells were fixed for 15 min with 4% PFA and washed with DPBS. KSHV(-) iSLK cells were washed in DPBS without fixing. After washing, cells were resuspended in DPBS. APC-A signal (for JF 646) was quantified by flow cytometry on a BD Bioscience LSR Fortessa. 10,000 events were collected per sample, and replicates represent independent reactivations on separate days. All data were analyzed using FlowJo software (BD Bioscience).

### Apoptosis and necrosis measurements

TRExRTA BCBL-1s with either empty vector, TFIIB-2xStrep, or TFIIB(D207A)-2xStrep were seeded at 1x10^6^ cells/mL and untreated or treated with 25 μM etoposide for 24 h. Cells were then labelled and analyzed by flow as described in a protocol from Crowley et al. (38) Briefly, cells were collected by centrifugation and resuspended in Annexin staining buffer (10 mM HEPES pH 7.4, 150 mM NaCl, 2.5 mM CaCl_2_, in DPBS (GIBCO)) with 1:10 dilution of Annexin V-FITC (fluorescein isothiocyanate; BD Biosciences) and 50 μg/mL propidium iodide (Sigma-Aldrich) for 15 minutes in the dark. Signal was quantified by flow cytometry on BD Bioscience LSR Fortessa using the FITC channel for Annexin-FITC, and PERCP channel for propidium iodide. 10,000 events were collected per sample and data were analyzed using FlowJo software (BD Bioscience) including color compensation. Small fragments were excluded from analysis, but gates were set to include both live and dead cells. Quadrants were set based on no dye and single dye controls and the percent population was determined for each quadrant. Ratios of population percentage normalized to empty vector were quantified.

### Data visualization and analysis

Data quantifications and statistical analysis were conducted in GraphPad Prism (v. 9-10.10). Statistical tests and significance values are noted in figure captions. The graphical abstract was generated using BioRender.com.

## Acknowledgements

We thank all members of the Glaunsinger and Coscoy labs (UC Berkeley) for their helpful feedback and comments on the manuscript, especially Azra Lari for assistance with imaging. This research was supported by NIH grants AI122528 and CA136367 to B.G., who is also an investigator of the Howard Hughes Medical Institute. We additionally thank the UC Davis Bioinformatics Core (RRID:SCR_023887) and the UC Berkeley QB3 Genomic Facilities (RRID:SCR_022170) for their services.

## Author contributions

Conceptualization, L.G. and B.A.G.; methodology, L.G. and B.A.G.; experimentation, L.G.; writing, L.G. and B.A.G.; formal analysis, L.G. and B.A.G.; project administration, L.G. and B.A.G; supervision and funding acquisition, B.A.G.

## Notes

### Competing Interest Statement

The authors have declared no competing interest.

### Summary of Updates

This revision includes minor text and citation modifications

## References

1. A. Vihervaara, F. M. Duarte, J. T. Lis, Molecular mechanisms driving transcriptional stress responses. Nature Reviews Genetics 19, 385–397 (2018).

2. M. F. Lavin, N. Gueven, The complexity of p53 stabilization and activation. Cell Death & Differentiation 13, 941–950 (2006).

3. L. Chen, S. Liu, Y. Tao, Regulating tumor suppressor genes: post-translational modifications. Signal Transduction and Targeted Therapy 5 (2020).

4. A. Tufegdžić Vidaković et al., Regulation of the RNAPII Pool Is Integral to the DNA Damage Response. Cell 180, 1245–1261.e1221 (2020).

5. C. Duncan-Lewis, E. Hartenian, V. King, B. A. Glaunsinger, Cytoplasmic mRNA decay represses RNA polymerase II transcription during early apoptosis. Elife 10 (2021).

6. N. A. Rosa-Mercado et al., Hyperosmotic stress alters the RNA polymerase II interactome and induces readthrough transcription despite widespread transcriptional repression. Mol Cell 81, 502–513.e504 (2021).

7. E. Hartenian, S. Gilbertson, J. D. Federspiel, I. M. Cristea, B. A. Glaunsinger, RNA decay during gammaherpesvirus infection reduces RNA polymerase II occupancy of host promoters but spares viral promoters. PLoS Pathog 16, e1008269 (2020).

8. L. Gulyas, B. A. Glaunsinger, RNA polymerase II subunit modulation during viral infection and cellular stress. Current Opinion in Virology 56, 101259 (2022).

9. E. Abernathy, S. Gilbertson, R. Alla, B. Glaunsinger, Viral Nucleases Induce an mRNA Degradation-Transcription Feedback Loop in Mammalian Cells. Cell Host Microbe 18, 243–253 (2015).

10. D. L. V. Bauer et al., Influenza Virus Mounts a Two-Pronged Attack on Host RNA Polymerase II Transcription. Cell Rep 23, 2119–2129.e2113 (2018).

11. K. A. Spriggs, M. Bushell, A. E. Willis, Translational Regulation of Gene Expression during Conditions of Cell Stress. Molecular Cell 40, 228–237 (2010).

12. M. Holcik, N. Sonenberg, Translational control in stress and apoptosis. Nature Reviews Molecular Cell Biology 6, 318–327 (2005).

13. B. Rozman, T. Fisher, N. Stern-Ginossar, Translation-A tug of war during viral infection. Mol Cell 83, 481–495 (2023).

14. S. Covarrubias, J. M. Richner, K. Clyde, Y. J. Lee, B. A. Glaunsinger, Host shutoff is a conserved phenotype of gammaherpesvirus infection and is orchestrated exclusively from the cytoplasm. J Virol 83, 9554–9566 (2009).

15. B. Glaunsinger, D. Ganem, Lytic KSHV infection inhibits host gene expression by accelerating global mRNA turnover. Mol Cell 13, 713–723 (2004).

16. C. P. Chen et al., Kaposi’s Sarcoma-Associated Herpesvirus Hijacks RNA Polymerase II To Create a Viral Transcriptional Factory. J Virol 91 (2017).

17. W. Deng, S. G. Roberts, TFIIB and the regulation of transcription by RNA polymerase II. Chromosoma 116, 417–429 (2007).

18. D. Kostrewa et al., RNA polymerase II–TFIIB structure and mechanism of transcription initiation. Nature 462, 323–330 (2009).

19. S. Sainsbury, J. Niesser, P. Cramer, Structure and function of the initially transcribing RNA polymerase II–TFIIB complex. Nature 493, 437–440 (2013).

20. B. N. Singh, M. Hampsey, A Transcription-Independent Role for TFIIB in Gene Looping. Molecular Cell 27, 806–816 (2007).

21. M. J. O’Brien, A. Ansari, Beyond the canonical role of TFIIB in eukaryotic transcription. Curr Genet 68, 61–67 (2022).

22. M. J. O’Brien, A. Ansari, Protein interaction network revealed by quantitative proteomic analysis links TFIIB to multiple aspects of the transcription cycle. Biochim Biophys Acta Proteins Proteom 1872, 140968 (2023).

23. J. F. Santana, G. S. Collins, M. Parida, D. S. Luse, D. H. Price, Differential dependencies of human RNA polymerase II promoters on TBP, TAF1, TFIIB and XPB. Nucleic Acids Res 50, 9127–9148 (2022).

24. J. Shandilya, Y. Wang, S. G. Roberts, TFIIB dephosphorylation links transcription inhibition with the p53-dependent DNA damage response. Proc Natl Acad Sci U S A 109, 18797–18802 (2012).

25. Y. Wang, J. A. Fairley, S. G. Roberts, Phosphorylation of TFIIB links transcription initiation and termination. Curr Biol 20, 548–553 (2010).

26. H. Nakamura et al., Global Changes in Kaposi’s Sarcoma-Associated Virus Gene Expression Patterns following Expression of a Tetracycline-Inducible Rta Transactivator. Journal of Virology 77, 4205–4220 (2003).

27. H. Wang, O. Julien, CaspSites: A Database and Web Application for Experimentally Observed Human Caspase Substrates Using N-Terminomics. Journal of Proteome Research 22, 454–461 (2023).

28. G. Y. Liu et al., Antizyme, a natural ornithine decarboxylase inhibitor, induces apoptosis of haematopoietic cells through mitochondrial membrane depolarization and caspases’ cascade. Apoptosis 11, 1773–1788 (2006).

29. P. L. Zhou et al., Perilipin 5 deficiency promotes atherosclerosis progression through accelerating inflammation, apoptosis, and oxidative stress. Journal of Cellular Biochemistry 120, 19107–19123 (2019).

30. J. Feng et al., Perilipin 5 ameliorates high-glucose-induced podocyte injury via Akt/GSK-3β/Nrf2-mediated suppression of apoptosis, oxidative stress, and inflammation. Biochem Biophys Res Commun 544, 22–30 (2021).

31. J. Liang et al., Mitochondrial PKM2 regulates oxidative stress-induced apoptosis by stabilizing Bcl2. Cell Research 27, 329–351 (2017).

32. L. Wang et al., Hypoxia-induced LncRNA DACT3-AS1 upregulates PKM2 to promote metastasis in hepatocellular carcinoma through the HDAC2/FOXA3 pathway. Experimental & Molecular Medicine 54, 848–860 (2022).

33. Q. Li, et al., Disruption of Robo2-Baiap2 integrated signaling drives cystic disease. JCI Insight 4 (2019).

34. S. Qian et al., The role of BCL-2 family proteins in regulating apoptosis and cancer therapy. Frontiers in Oncology 12 (2022).

35. Y. Wang et al., ALPK2 acts as tumor promotor in development of bladder cancer through targeting DEPDC1A. Cell Death & Disease 12 (2021).

36. C. Meneses et al., The angiotensin-(1–7)/Mas axis reduces myonuclear apoptosis during recovery from angiotensin II-induced skeletal muscle atrophy in mice. Pflügers Archiv - European Journal of Physiology 467, 1975–1984 (2015).

37. L. Zhang, M. Yao, W. Ma, Y. Jiang, W. Wang, MicroRNA-376b-3p targets RGS1 mRNA to inhibit proliferation, metastasis, and apoptosis in osteosarcoma. Annals of Translational Medicine 9, 1652–1652 (2021).

38. L. C. Crowley, B. J. Marfell, A. P. Scott, N. J. Waterhouse, Quantitation of Apoptosis and Necrosis by Annexin V Binding, Propidium Iodide Uptake, and Flow Cytometry. Cold Spring Harbor Protocols 2016, pdb.prot087288 (2016).

39. M. Watanabe et al., A substrate-trapping strategy to find E3 ubiquitin ligase substrates identifies Parkin and TRIM28 targets. Communications Biology 3 (2020).

40. G. Perel, J. Bliss, C. M. Thomas, Carfilzomib (Kyprolis): A Novel Proteasome Inhibitor for Relapsed And/or Refractory Multiple Myeloma. P t 41, 303–307 (2016).

41. M. L. Hyer et al., A small-molecule inhibitor of the ubiquitin activating enzyme for cancer treatment. Nature Medicine 24, 186–193 (2018).

42. B. Herquel et al., Transcription cofactors TRIM24, TRIM28, and TRIM33 associate to form regulatory complexes that suppress murine hepatocellular carcinoma. Proceedings of the National Academy of Sciences 108, 8212–8217 (2011).

43. V. M. Advani, P. Ivanov, Translational Control under Stress: Reshaping the Translatome. BioEssays 41, 1900009 (2019).

44. M. D. Wilson, M. Harreman, J. Q. Svejstrup, Ubiquitylation and degradation of elongating RNA polymerase II: The last resort. Biochimica et Biophysica Acta (BBA) - Gene Regulatory Mechanisms 1829, 151–157 (2013).

45. P. Caron et al., WWP2 ubiquitylates RNA polymerase II for DNA-PK-dependent transcription arrest and repair at DNA breaks. Genes & Development 33, 684–704 (2019).

46. M. S. D’Arcy, Cell death: a review of the major forms of apoptosis, necrosis and autophagy. Cell Biology International 43, 582–592 (2019).

47. S. R. Rosario et al., Pan-cancer analysis of transcriptional metabolic dysregulation using The Cancer Genome Atlas. Nature Communications 9 (2018).

48. S. Shiraishi, N. Tamamura, M. Jogo, Y. Tanaka, T.-A. Tamura, Rapid proteasomal degradation of transcription factor IIB in accordance with F9 cell differentiation. Gene 436, 115–120 (2009).

49. M. J. O’Brien, A. Ansari, Critical Involvement of TFIIB in Viral Pathogenesis. Front Mol Biosci 8, 669044 (2021).

50. D. A. Haas et al., Viral targeting of TFIIB impairs de novo polymerase II recruitment and affects antiviral immunity. PLOS Pathogens 14, e1006980 (2018).

51. C. Vogt et al., The interferon antagonist ML protein of thogoto virus targets general transcription factor IIB. J Virol 82, 11446–11453 (2008).

52. R. Gupta et al., Characterization of the interaction between the acidic activation domain of VP16 and the RNA polymerase II initiation factor TFIIB. Nucleic Acids Res 24, 2324–2330 (1996).

53. F. Hayashi et al., Human General Transcription Factor TFIIB: Conformational Variability and Interaction with VP16 Activation Domain. Biochemistry 37, 7941–7951 (1998).

54. R. Caswell et al., The human cytomegalovirus 86K immediate early (IE) 2 protein requires the basic region of the TATA-box binding protein (TBP) for binding, and interacts with TBP and transcription factor TFIIB via regions of IE2 required for transcriptional regulation. Journal of General Virology 74, 2691–2698 (1993).

55. X. Tong, F. Wang, C. J. Thut, E. Kieff, The Epstein-Barr virus nuclear protein 2 acidic domain can interact with TFIIB, TAF40, and RPA70 but not with TATA-binding protein. Journal of Virology 69, 585–588 (1995).

56. V. Gelev et al., A new paradigm for transcription factor TFIIB functionality. Scientific Reports 4 (2014).

57. Zoe et al., Global Mapping of Herpesvirus-Host Protein Complexes Reveals a Transcription Strategy for Late Genes. Molecular Cell 57, 349–360 (2015).

58. K. Randolph, U. Hyder, I. D’Orso, KAP1/TRIM28: Transcriptional Activator and/or Repressor of Viral and Cellular Programs? Frontiers in Cellular and Infection Microbiology 12 (2022).

59. P.-C. Chang et al., Kruppel-Associated Box Domain-Associated Protein-1 as a Latency Regulator for Kaposi’s Sarcoma-Associated Herpesvirus and Its Modulation by the Viral Protein Kinase. Cancer Research 69, 5681–5689 (2009).

60. C. A. King, Kaposi’s Sarcoma-Associated Herpesvirus Kaposin B Induces Unique Monophosphorylation of STAT3 at Serine 727 and MK2-Mediated Inactivation of the STAT3 Transcriptional Repressor TRIM28. Journal of Virology 87, 8779–8791 (2013).

61. X. Li, E. M. Burton, S. Bhaduri-Mcintosh, Chloroquine triggers Epstein-Barr virus replication through phosphorylation of KAP1/TRIM28 in Burkitt lymphoma cells. PLOS Pathogens 13, e1006249 (2017).

62. X. Li, S. V. Kozlov, A. El-Guindy, S. Bhaduri-Mcintosh, Retrograde Regulation by the Viral Protein Kinase Epigenetically Sustains the Epstein-Barr Virus Latency-to-Lytic Switch To Augment Virus Production. Journal of Virology 93 (2019).

63. H. Xu, I. A. Akinyemi, J. Haley, M. T. McIntosh, S. Bhaduri-Mcintosh, ATM, KAP1 and the Epstein–Barr virus polymerase processivity factor direct traffic at the intersection of transcription and replication. Nucleic Acids Research 51, 11104–11122 (2023).

64. C. F. De La Cruz-Herrera, K. Shire, U. Z. Siddiqi, L. Frappier, A genome-wide screen of Epstein-Barr virus proteins that modulate host SUMOylation identifies a SUMO E3 ligase conserved in herpesviruses. PLOS Pathogens 14, e1007176 (2018).

65. D. Silva, G. Santos, M. Barroca, D. Costa, T. Collins, “Inverse PCR for Site-Directed Mutagenesis”. (Springer US, 2023), 10.1007/978-1-0716-3358-8_18, pp. 223–238.

66. E. Kowarz, D. Löscher, R. Marschalek, Optimized Sleeping Beauty transposons rapidly generate stable transgenic cell lines. Biotechnology Journal 10, 647–653 (2015).

67. K. F. Brulois et al., Construction and Manipulation of a New Kaposi’s Sarcoma-Associated Herpesvirus Bacterial Artificial Chromosome Clone. Journal of Virology 86, 9708–9720 (2012).

68. J. Myoung, D. Ganem, Generation of a doxycycline-inducible KSHV producer cell line of endothelial origin: Maintenance of tight latency with efficient reactivation upon induction. Journal of Virological Methods 174, 12–21 (2011).

69. M. Stürzl, D. Gaus, W. G. Dirks, D. Ganem, R. Jochmann, Kaposi’s sarcoma-derived cell line SLK is not of endothelial origin, but is a contaminant from a known renal carcinoma cell line. International Journal of Cancer 132, 1954–1958 (2013).

70. N. Labo et al., Heterogeneity and Breadth of Host Antibody Response to KSHV Infection Demonstrated by Systematic Analysis of the KSHV Proteome. PLoS Pathogens 10, e1004046 (2014).

71. E. Hartenian, A. S. Mendez, A. L. Didychuk, S. Khosla, Britt, DNA processing by the Kaposi’s sarcoma-associated herpesvirus alkaline exonuclease SOX contributes to viral gene expression and infectious virion production. Nucleic Acids Research 51, 182–197 (2023).

72. L. S. Toni et al., Optimization of phenol-chloroform RNA extraction. MethodsX 5, 599–608 (2018).

73. J. Ohata, F. Vohidov, Z. T. Ball, Convenient analysis of protein modification by chemical blotting with fluorogenic “click” reagents. Molecular BioSystems 11, 2846–2849 (2015).

74. J. Schindelin et al., Fiji: an open-source platform for biological-image analysis. Nature Methods 9, 676–682 (2012).

75. A. Dobin et al., STAR: ultrafast universal RNA-seq aligner. Bioinformatics 29, 15–21 (2013).

76. C. W. Law et al., RNA-seq analysis is easy as 1-2-3 with limma, Glimma and edgeR. F1000Research 5, 1408 (2018).

