## Supplemental information for "The general transcription factor TFIIB is a target for transcriptome control during cellular stress and viral infection" (from another version of this preprint: version 2, 2024-05-03)

**This PDF file includes:**

Figures S1 to S6

Tables S1 to S2

SI References

#### Supplemental Figures

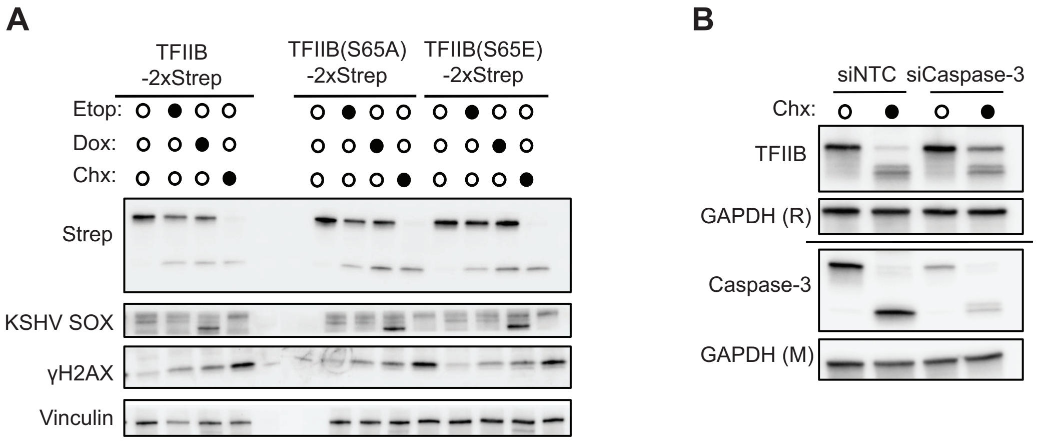

**Figure S1. Caspase-3 cleaves TFIIB independent of the TFIIB phosphorylation state. (A)** TRExRTA BCBL-1 cells expressing TFIIB-2xStrep, TFIIB(S65A)-2xStrep, or TFIIB(S65E)-2xStrep exposed to 25 μM etoposide (Etop; DNA damage agent) for 24 h, 2 μg/mL doxycycline (Dox; viral reactivation) for 24h, or 100 μg/mL cycloheximide (Chx; translational inhibitor) for 6 h, or mock treated with drug diluent and analyzed by western blotting. SOX is a marker for viral infection and γH2AX is induced during DNA damage. **(B)** Caspase-3 was depleted by siRNA in TRExRTA BCBL-1 for 48h followed by 4h of 100 μg/mL Chx and analyzed by western blotting; GAPDH probed with rabbit antibody is a loading control for TFIIB, and GAPDH probed with mouse antibody is a loading control for caspase-3.

**Figure S1**

**Figure S1. Caspase-3 cleaves TFIIB independent of the TFIIB phosphorylation state. (A)** TRExRTA BCBL-1 cells expressing TFIIB-2xStrep, TFIIB(S65A)-2xStrep, or TFIIB(S65E)-2xStrep exposed to 25 μM etoposide (Etop; DNA damage agent) for 24 h, 2 μg/mL doxycycline (Dox; viral reactivation) for 24h, or 100 μg/mL cycloheximide (Chx; translational inhibitor) for 6 h, or mock treated with drug diluent and analyzed by western blotting. SOX is a marker for viral infection and γH2AX is induced during DNA damage. **(B)** Caspase-3 was depleted by siRNA in TRExRTA BCBL-1 for 48h followed by 4h of 100 μg/mL Chx and analyzed by western blotting; GAPDH probed with rabbit antibody is a loading control for TFIIB, and GAPDH probed with mouse antibody is a loading control for caspase-3.

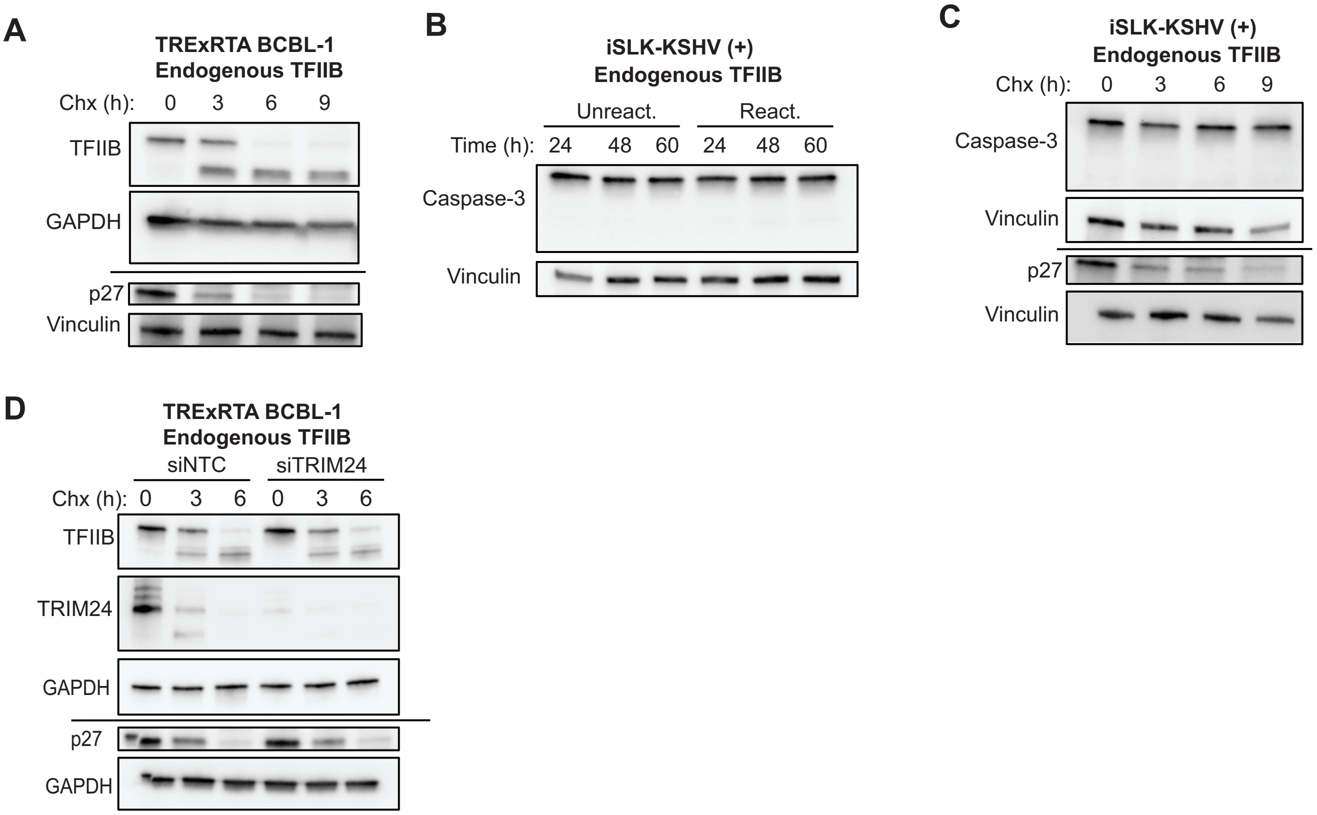

**Figure S2. Control blots supporting caspase-independent turnover of TFIIB in iSLK cells and showing that TRIM24 does not influence TFIIB stabilization. (A)** TRExRTA BCBL-1 cells treated with 100 μg/mL Chx and endogenous TFIIB protein were analyzed by western blotting. p27 is a control for translational inhibition and GAPDH probed with mouse antibody and vinculin are loading controls. **(B)** iSLK-KSHV(+) BAC16 cells were reactivated with doxycycline for the given timepoints prior to western blotting. Vinculin serves as a loading control. Note the lack of appreciable procaspase-3 cleavage and activation. These samples come from the same experiment as *Fig. 3C*. **(C)** iSLK-KSHV(+) BAC16 cells treated with 250 μg/mL cycloheximide (Chx) for given timepoints. Note lack of procaspase-3 activation. **(D)** Western blots from of TRExRTA BCBL-1 cells nucleofected with TRIM24 siRNA or non-targeting control (siNTC) for 18 h followed by a timecourse with 100 μg/mL cycloheximide treatment (Chx). p27 is a translation inhibitor control and GAPDH probed with mouse antibody a loading control.

**Figure S2**

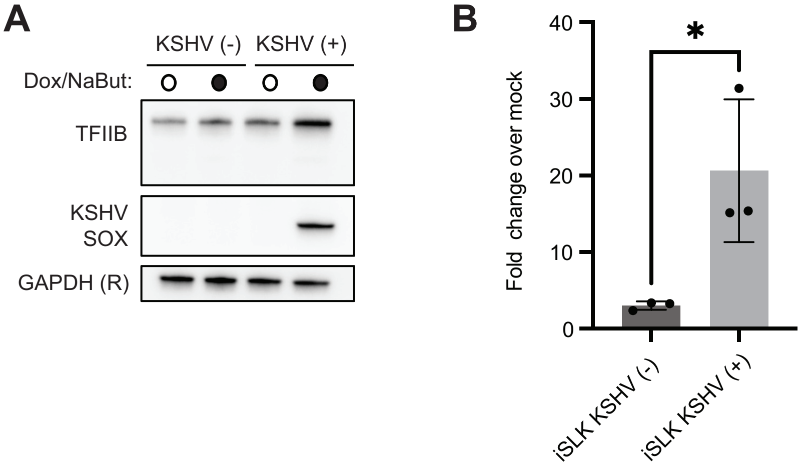

**Figure S3. TFIIB stabilization requires KSHV lytic reactivation and is not induced in unreactivated cells or iSLK cells lacking virus. (A)** Western blotting of iSLK cells containing or lacking the KSHV BAC16. Cells were mock treated or dosed with 5 ug/mL doxycycline (Dox) and 1 mM sodium butyrate (NaBut) for 48 h. KSHV SOX is a marker for lytic viral reactivation and anti-GAPDH probed with rabbit antibody is a loading control. Flow analysis of iSLK cells expressing TFIIB-Halo and containing or lacking the KSHV BAC16. (**B**) Flow analysis of iSLK cells expressing TFIIB-Halo and containing or lacking the KSHV BAC16. Cells were treated with Dox and NaBut and harvested 48h later. Halo was labeled with JF646-Halo dye and was measured by the signal in the APC-A channel and the fold change in fluorescence over mock-treated cells was quantified. *$P < 0.0307$; Student's unpaired t-test.

**Figure S3**

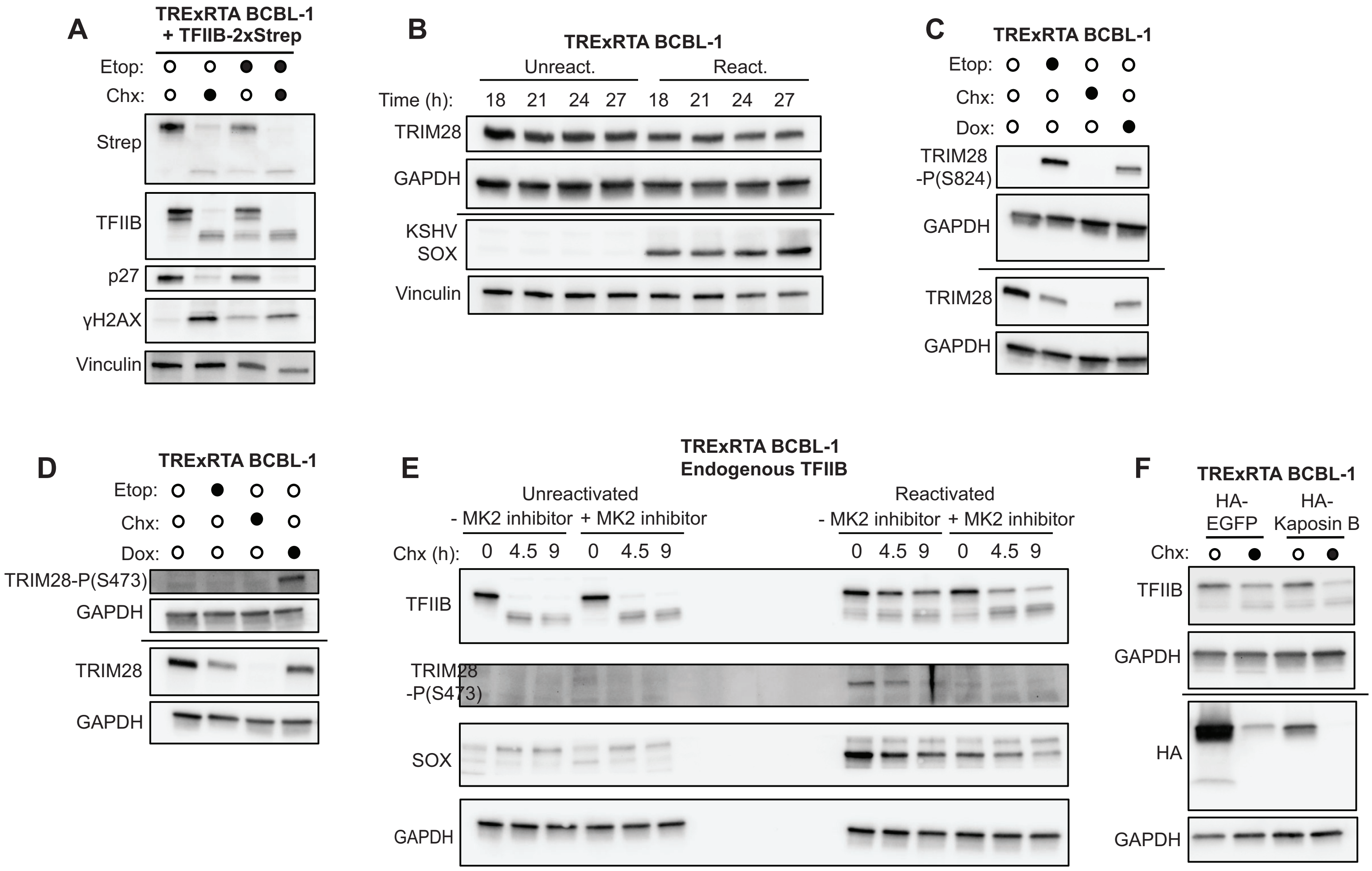

A
TRExRTA BCBL-1
+ TFIIB-2xStrep
Etop:
Chx:
Strep
TFIIB
p27
γH2AX
Vinculin
B
TRExRTA BCBL-1
Unreact.
React.
Time (h):
18 21 24 27 18 21 24 27
TRIM28
GAPDH
KSHV
SOX
Vinculin
C
TRExRTA BCBL-1
Etop:
Chx:
Dox:
TRIM28
-P(S824)
GAPDH
TRIM28
GAPDH
D
TRExRTA BCBL-1
Etop:
Chx:
Dox:
TRIM28-P(S473)
GAPDH
TRIM28
GAPDH
E
TRExRTA BCBL-1
Endogenous TFIIB
Unreactivated
Reactivated
- MK2 inhibitor
+ MK2 inhibitor
- MK2 inhibitor
+ MK2 inhibitor
Chx (h):
0 4.5 9 0 4.5 9 0 4.5 9 0 4.5 9
TFIIB
TRIM28
-P(S473)
SOX
GAPDH
F
TRExRTA BCBL-1
HA-
EGFP
HA-
Kaposin B
Chx:
TFIIB
GAPDH
HA
GAPDH

Figure S4

**Figure S4. Neither TRIM28 protein levels nor specific phosphorylation events account for KSHV-induced TFIIB stabilization. (A)** DNA damage does not induce TFIIB stabilization. TRExRTA BCBL-1 cells expressing TFIIB-2xStrep were exposed to 25 μM etoposide (Etop; DNA damage agent) or mock treatment for 16 h followed by 6 h of 100 μg/mL cycloheximide (Chx) to stall translation. **(B)** KSHV reactivation does not cause TFIIB stabilization by altering TRIM28 levels. Western analysis of endogenous TRIM28 in TRExRTA BCBL-1 cells over a timecourse post-KSHV reactivation or mock treatment. KSHV SOX is a marker for reactivation, and vinculin and GAPDH probed with mouse antibody are loading controls.

**(C-D)** Two major TRIM28 phosphoregulatory events occur at its S473 and S824 residues (1, 2). To determine whether either modification occurs during stress, TRExRTA BCBL-1 cells were treated with 25 μM etoposide (Etop; DNA damage agent) for 24 h, 2 μg/mL doxycycline (Dox; viral reactivation) for 24h, or 100 μg/mL cycloheximide (Chx; translational inhibitor) for 6 h, or mock treated with drug diluent and analyzed by western blotting. GAPDH probed with mouse antibody serves as a loading control. (*C*) shows that TRIM28-P(S824) is present in both etoposide-treated or reactivated cells and is thus unlikely to be responsible for TFIIB stabilization since etoposide treatment does not stabilize TFIIB (*A*). (*D*) show that TRIM28-P(S473) is present only during viral reactivation. **(E-F)** KSHV gene Kaposin B activates MAPK-activated protein kinase 2 (MK2) to stimulate phosphorylation of TRIM28 S473 (3, 4). To test whether this could explain TFIIB stabilization, TRExRTA BCBL-1 cells +/- reactivation with dox were simultaneously treated with 20 μM of the MK2 inhibitor Gamcemetinib (CC-99677) or diluant control for 16 h followed by 9 h of 100 μg/mL cycloheximide (Chx) to stall translation. GAPDH probed with rabbit antibody serves as a loading control. At concentrations of MK2 inhibitor sufficient to prevent TRIM28-P(S473), the KSHV viral lifecycle was impaired as measured through protein levels of the KSHV early gene SOX. While TFIIB half-life was somewhat reduced, this is most likely a consequence of reduced KSHV reactivation rather than inhibition of TRIM28-P(S473). **(F)** To more directly monitor if KSHV Kaposin B expression stabilizes TFIIB, TRExRTA BCBL-1 cells were nucleofected with vectors driving either HA-tagged Kaposin B or a control HA-tagged EGFP of similar size for 18 h followed by 9 h of 100 μg/mL Chx. GAPDH probed with mouse antibody (top) and rabbit antibody (bottom) serve as loading controls. There was no change in stability during Kaposin B transfection relative to the EGFP control. Thus, neither MK2 inhibition nor Kaposin B transfection significantly altered TFIIB levels.

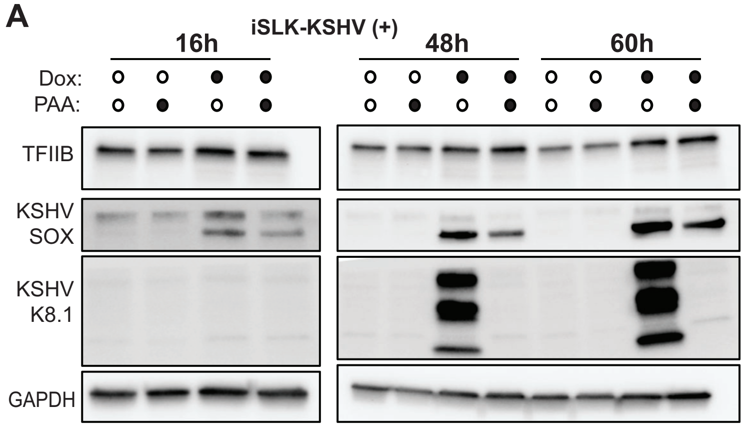

**Figure S5. TFIIB stabilization is an early event in reactivated iSLK cells.** Western blots of iSLK-KSHV(+) BAC16 cells +/- reactivation and treated with 1mM of phosphonoacetic acid (PAA). SOX is a viral early gene and K8.1 is a late gene (expressed only after DNA replication).

**Figure S5**

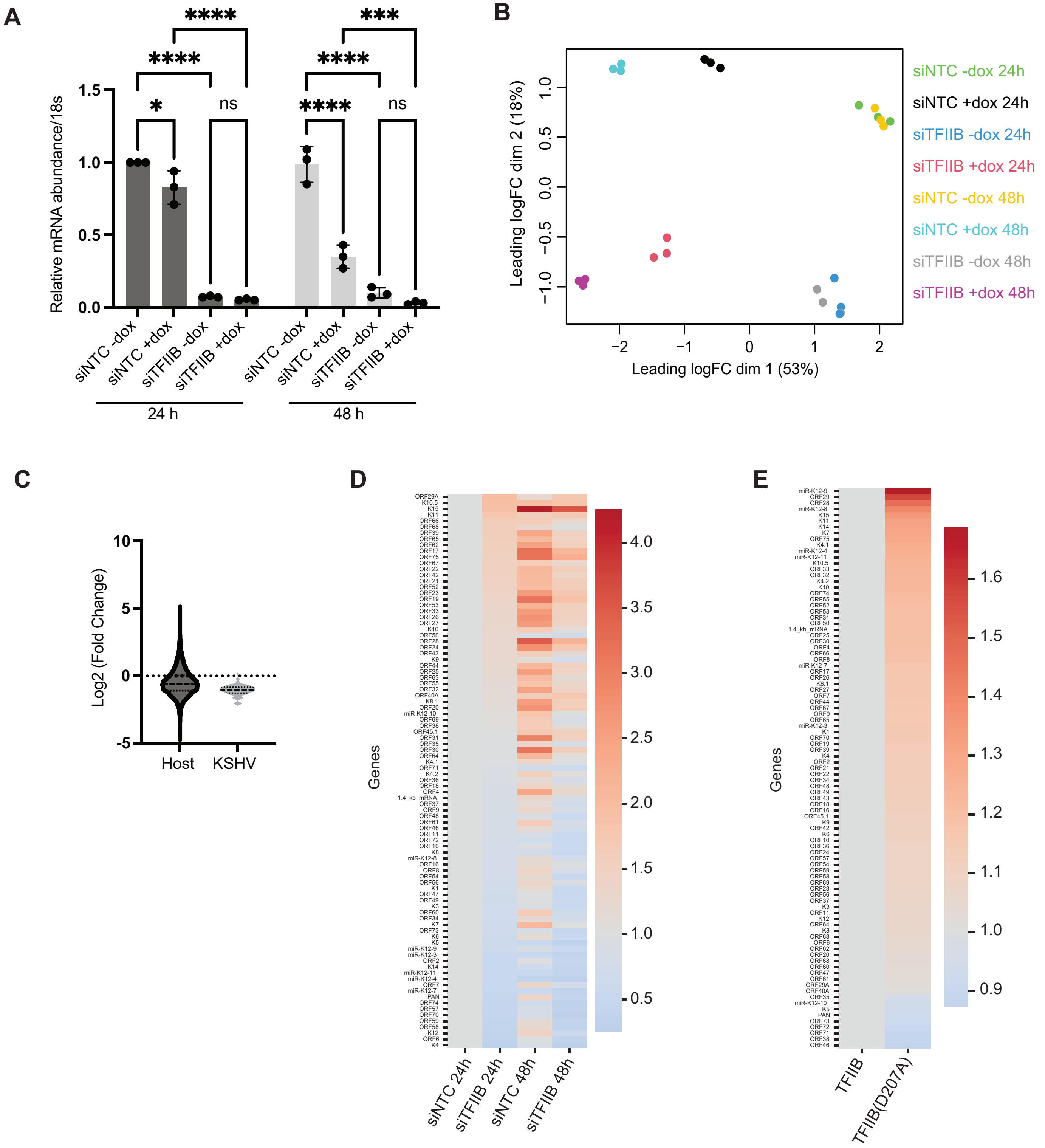
A
Relative mRNA abundance/18s
1.5
1.0
0.5
0.0
****
****
*
ns
***
****
****
ns
siNTC -dox
siNTC +dox
siTFIIB -dox
siTFIIB +dox
24 h
48 h
B
Leading logFC dim 2 (18%)
Leading logFC dim 1 (53%)
siNTC -dox 24h
siNTC +dox 24h
siTFIIB -dox 24h
siTFIIB +dox 24h
siNTC -dox 48h
siNTC +dox 48h
siTFIIB -dox 48h
siTFIIB +dox 48h
C
Log2 (Fold Change)
10
5
0
-5
Host
KSHV
D
Genes
siNTC 24h
siTFIIB 24h
siNTC 48h
siTFIIB 48h
4.0
3.5
3.0
2.5
2.0
1.5
1.0
0.5
E
Genes
TFIIB
TFIIB(D207A)
1.6
1.5
1.4
1.3
1.2
1.1
1.0
0.9

**Figure S6**

**Figure S6. TFIIB depletion impairs KSHV lytic gene expression. (A)** RT-qPCR of TFIIB RNA normalized to 18S rRNA to confirm TFIIB knockdown in TRExRTA BCBL-1 cells nucleofected with TFIIB siRNA (siTFIIB) or non-targeting siRNA (siNTC) followed by 24 h or 48 h of doxycycline (dox)-induced reactivation or mock. ns- nonsignificant, **** $P < 0.0001$, *** $P = 0.0004$, * $P = 0.0285$; two-way ANOVA with Tukey correction. **(B)** PCA analysis of RNA-seq samples described in (*A*) demonstrating triplicate consistency. **(C)** Violin plot of gene expression changes for differentially expressed cellular (host) and viral genes from RNA-seq analysis of siTFIIB knockdown relative to siNTC control at 48 h post-reactivation. This is the ensemble of all changes unfiltered by P value. ****$P < 0.0001$; Student's T-test with Welch correction. **(D)** Heatmap of KSHV viral gene expression changes in RNA-seq from siTFIIB and siNTC knockdown at the given timepoints. Cells were reactivated in all conditions and normalized to 24 h gene expression in siNTC cells. **(E)** Heatmap of viral gene expression changes from RNA-seq analysis of TRExRTA BCBL-1 cells expressing TFIIB-2xStrep or TFIIB(D207A)-2xStrep and reactivated for 48 h

**Supplemental Table 1.** Identities and roles of significantly upregulated apoptosis-associated genes in TFIIB(D207A) expressing cells during TRAIL treatment from RNA-seq analysis in main *Fig. 1F*.

| Gene Name | Function | Role in Apoptosis | Citation |
|---|---|---|---|
| **AZIN2** | Antizyme inhibitor 2 | Inhibitor of antizyme. Antizyme overexpression enhances apoptotic cell death and interferes with BCL-2 expression. | (5) Liu et al. 2006 |
| **PLIN5** | Perilipin 5 | Knockdown of PLIN5 in APO (-/-) mice leads is pro-apopotic and leads to caspase-3 activation (1). PLIN5 overexpression supresses apoptosis in high glucose-injured podocytes (2). | (6) Zhou et al. 2019 (7) Feng et al. 2021 |
| **DACT3-AS1** | DACT3 antisense RNA 1 | (1-2) Stabilizes PMK2 when induced by hypoxia. PMK2 stabilizes Bcl1 to supress apoptosis. (3) Note, though, that DACT3-AS1 loss also appears to be a positive regulator of ferroptosis. | (8)Wang et al. 2022 (9) Liang et al. 2017 (10) Qu et al. 2022 |
| **BAIAP2** | BAR/IMD domain containing adaptor protein 2 | Forms a dimer with Robo2 to promote MDM2 phosphorylation and degradation of p53. Knockdown upregulates DNA damage and p53 levels, leading to senescence. | (11) Li et al. 2019 |
| **BCL2** | B-cell lymphoma 2 | Heterodimerizes with BAX and regulates calcium levels for pro-survival effect. Prevents activities of some caspases. | (12) Qian et al. 2022 |
| **ALPK2** | Alpha kinase 2 | ALPK2 knockdown promotes cell apoptosis in studies of badder cancer. ALPK2 knockdown upregulates a suite of apoptosis-related genes. | (13) Wang et al. 2021 |
| **MAS1** | MAS1 proto-oncogene, G protein-coupled receptor | MAS1 interacts with angiotensin-(1-7) and decreases apoptotic events in skeletal muscle cess during recovery from atrophy. | (14) Meneses et al. 2015 |
| **RGS1** | Regulator of G protein signaling 1 | RGS1 has a proliferative effect on osteosarcoma cells. Treating cells with miR-376b-3p, which targets RGS1, leads to increased apoptosis. | (15) Zhang et al. 2021 |

**Supplemental Table 2.** Materials and reagents used in this study.

| Reagent or resource | Source | Identifier/citation |
|---|---|---|
| ***Antibodies*** | | |
| **Mouse anti- TFIIB (1:1000), clone 2F6A3H4** | Cell Signaling | 4169 |
| **Mouse anti-GAPDH (1:5000), clone 6C5** | Thermofisher | AM4300 |
| **Rabbit anti-GAPDH (1:1000), clone 14C10** | Cell Signaling | 2118 |
| **Rabbit anti-p27 (1:1000), polyclonal** | ProteinTech | 15086-1-AP |
| **Rabbit anti-TRIM28 (1:1000), polyclonal** | Cell Signaling | 4123 |
| **Rabbit anti-TRIM28 -P(S824) (1:1000), polyclonal** | Cell Signaling | 4127 |
| **Rabbit anti-TRIM28-P(S473) (1:500-1:1000), clone 11G10SC** | BioLegend | 654101 |
| **Rabbit anti-HA (1:1000), clone C29F4** | Cell Signaling | 3742 |
| **Rabbit anti-TRIM24 (1:1000), clone E93TN** | Cell Signaling | 79030 |
| **Rabbit anti-Vinculin (1:1000), polyclonal** | Abcam | ab91459 |
| **Mouse anti-FK2 (1:1000), polyclonal** | Enzo | BML-PW8810-0100 |
| **Rabbit anti-NRF2 (1:1000), polyclonal** | ProteinTech | 16396-1-AP |
| **Rabbit anti-Caspase-3 (1:1000), polyclonal** | Cell Signaling | 9662 |
| **Rabbit anti-Caspase-8 (1:1000), clone E7** | Abcam | ab32397 |
| **Rabbit anti-γH2AX (1:1000), polyclonal** | Bethyl | A300-081A-M |

| Reagent or resource | Source | Identifier/citation |
|---|---|---|
| **Rabbit anti-p53 (1:1000), clone 1C12** | Cell Signaling | 2524 |
| **Goat Anti-Mouse IgG(H+L) Human ads-HRP (1:5000)** | Southern Biotech | 100242-504 |
| **Goat Anti-Rabbit IgG(H+L) Human ads-HRP (1:5000)** | Southern Biotech | 4030-05 |
| **Strep-Tag II Antibody HRP Conjugate (1:2000)** | Sigma Aldrich | 71591-3 |
| **Rabbit anti-K8.1 (1:10000)** | Gift of Denise Whitby | Labo et al. 2014 (16) |
| **Rabbit anti-SOX (1:5000)** | Lab stock; Yenzyme | Hartenian et al. 2023 (17), YZ6540 (cycle 1) |
| ***Human cell lines and virus information*** | | |
| **iSLK-puromycin (doxycylcine-inducible RTA cassette)** | Lab stock | Brulois et al. 2012 (18); Myoung 2011(19); Stürzl 2013 (20) |
| **iSLK-puromycin (doxycylcine-inducible RTA cassette) + Halo-TFIIB** | This study | N/A |
| **iSLK + KSHV BAC16 (doxycyline-inducble RTA cassette and hygromycin-resistant viral BAC)** | Lab stock | Brulois et al. 2012 (18); Myoung 2011(19); Stürzl 2013 (20) |
| **iSLK + KSHV BAC16 (doxycyline-inducble RTA cassette and hygromycin-resistant viral BAC) + Halo** | This study | N/A |

| Reagent or resource | Source | Identifier/citation |
| --- | --- | --- |
| **iSLK + KSHV BAC16 (doxycyline-inducble RTA cassette and hygromycin-resistant viral BAC) + Halo-TFIIB** | This study | N/A |
| **Lenti-X 293T cells** | Clonetech/Takara | 632180 |
| **TRExRTA BCBL-1** | Gift of Jae Jung | Nakamura et al. 2003 (21) |
| **TRExRTA BCBL-1 + TFIIB-2xStrep** | This study | N/A |
| **TRExRTA BCBL-1 + TFIIB(D207A)-2xStrep** | This study | N/A |
| **TRExRTA BCBL-1 + TFIIB(S65A)-2xStrep** | This study | N/A |
| **TRExRTA BCBL-1 + TFIIB(S65E)-2xStrep** | This study | N/A |
| **Chemicals and other reagents** | | |
| ***Tissue Culture*** | | |
| **1X Glutamax** | Gibco | 35050061 |
| **RPMI 1640 Medium** | Gibco | 11875119 |
| **DMEM** | Gibco | 11965-118 |
| **DPBS 1x without Ca, Mg** | Gibco | 14190-250 |
| **Neon Transfection System, 100 uL Kit** | Invitrogen | MPK10096 |
| **On-Target Plus Non-targeting control pool** | Dharmacon | D-001810-10 |

| Reagent or resource | Source | Identifier/citation |
|---|---|---|
| **Accell Non-targeting Control Pool** | Dharmacon | D-001910-10 |
| **GTF2B ON-TARGETplus siRNA, SmartPool** | Dharmacon | L-012964-00 |
| **Accell Human GTF2B siRNA, SMARTPool** | Dharmacon | E-012964-00 |
| **ON-TARGETplus Human TRIM24 siRNA, SmartPool** | Dharmacon | L-005387-00 |
| **Accell Human TRIM28 siRNA, SMARTPool** | Dharmacon | E-005046-00 |
| **Accell Human CASP3 siRNA, SmartPool** | Dharmacon | E-004307-00-0005 |
| **ON-TARGETplus Human CASP3 siRNA, SmartPool** | Dharmacon | L-004307-00-0005 |
| ***Trans*IT-X2 Transfection Reagent** | Mirus | MIR 6003 |
| ***Trans*IT-LT1 Transfection Reagent** | Mirus | MIR 2300 |
| **SuperKillerTRAIL** | Enzo | ALX-201-115-C010 |
| **SUPERKILLERTRAIL Storage and Dilution Buffer** | Enzo | ENZ-PRT317-0500 |
| **Cycloheximide (solution in DMSO)** | Sigma | C4859-1ML |
| **Click-IT L-Homopropargylglycine (HPG)** | Thermo Scientific/Fisher | C10186 |
| **Janelia Fluor 646 HaloTag** | Promega | GA1121 |
| **BD Pharmingen™ FITC Annexin V (100 assays)** | BD Biosciences | 556420 |
| **Z-FA-FMK (DMSO solution)** | Santa Cruz | sc-201303 |
| **Z-VAD(OMe)-FMK (DMSO solution)** | Santa Cruz | sc-311561A |
| **Z-DEVD-FMK, Caspase-3 Inhibitor (DMSO solution)** | BD Sciences | 550378 |

| Reagent or resource | Source | Identifier/citation |
| --- | --- | --- |
| **Carfilzomib (PR-171) (DMSO solution)** | Selleckchem | 50-194-7948 |
| **TAK-234 (MLN7243) (DMSO solution)** | MedChemExpress via Cedarlane | HY-100487 |
| **Etoposide 10mM (DMSO solution)** | APExBio via Fisher | A1971 |
| **Phosphonoacetic acid (aqeous)** | Sigma | 284270-50G |
| **Tet-free Fetal Bovine Serum (FBS)** | UC Berkeley Tissue Culture Facility | N/A |
| **USDA-certified Fetal Bovine Serum (FBS)** | Peak Serum | PS-FB2 |
| **Gamcemetinib (CC-99677) (DMSO solution)** | Selleckchem | E1657 |
| ***Other standard laboratory drugs and chemicals were sourced from multiple vendors (e.g. antibiotics, etc.).** | | |
| ***Molecular biology*** | | |
| **In-Fusion HD Cloning Kit, 100 rxns** | Clonetech/Takara | 639650 |
| **Clarity Western ECL Substrate** | Bio-Rad | 1705061 |
| **Radiance plus AC2103 Femtogram-sensitivity HRP Substrate** | Azure Biosystems via VWR | 10147-298 (KT) |
| **Halt Protease and Phosphatase Inhibitor Cocktail (100X)** | Thermo Scientific/Fisher | PI78440 |
| **Benzonase Nuclease** | Millipore-Sigma | E1014-25KU |
| **Pierce 660nm Protein Assay Reagent** | Thermo Scientific/Fisher | 22660 |

| Reagent or resource | Source | Identifier/citation |
|---|---|---|
| **Ionic Detergent Compatibility Reagent for Pierce 660nm Protein Assay Reagent** | Thermo Scientific/Fisher | 22663 |
| **Laemmli Buffer, 2x** | Bio-Rad | 1610737 |
| **PageRuler Prestained Protein Ladder, 10 to 180 kDa** | Thermo Scientific/Fisher | 26620 |
| **Trans-Blot Turbo Transfer, Mini PVDF (example materials pack)** | Bio-Rad | 17001917 |
| **TRIzol** | Invitrogen | 15596018 |
| **Turbo DNase** | Thermo Scientific/Fisher | AM2238 |
| **AMV-RT** | Promega | M5108 |
| **iTaq UniverSYBR Green Supermix** | Bio-Rad | 1725125 |
| **QiaAMP Blood Mini kit** | Qiagen | 51104 |
| **MagStrep “type3” XT beads** | IBA Lifesciences | 2-4090-002 |
| **picolyl-azide-sulfo-Cy5** | Jena Biosciences | CLK-1177-1 |
| **Qubit 3 Fluorometer HS RNA kit** | Invitrogen | Q32855 |
| **ERCC RNA Spike-in mix** | Thermo Scientific/Fisher | 4456740 |
| **KAPA Unique Dual-Indexed Adapter Kit (15 µM) (virtual kit)-4x96 reactions** | Roche | 8861919702 |
| **KAPA Stranded RNA-seq Kit with RiboErase (HMR)** | Roche | KK8484 |

| Reagent or resource | Source | Identifier/citation |
|---|---|---|
| **KAPA RNA Hyperprep Kit with RiboErase** | Roche | KK8561 |
| **KAPA Library Quantification Kit** | Roche | KK4824 |
| ***Recombinant DNA/plasmids*** | | |
| **pcDNA4/TO-2xStrep (pCDNA4/TO Invitrogen vector modified to include a C-terminal 2xStrep tag)** | Invitrogen | V102020 |
| **pSBbi-GP (modified to remove EGFP)** | Addgene | #60511 |
| **pSBbi-GP (modified to remove EGFP)-TFIIB-2xStrep** | This study | Addgene ID pending |
| **pSBbi-GP (modified to remove EGFP)-TFIIB(D207A)-2xStrep** | This study | Addgene ID pending |
| **pSBbi-GP (modified to remove EGFP)-TFIIB(S65A)-2xStrep** | This study | Addgene ID pending |
| **pSBbi-GP (modified to remove EGFP)-TFIIB(S65E)-2xStrep** | This study | Addgene ID pending |
| **Sleeping beauty transposase** | Gift of Nicholas Ingolia | N/A |
| **pSBtet-Bla** | Addgene | #60510 |
| **pMD2.G** | Addgene | #12259 |
| **psPAX2** | Addgene | #12260 |

| Reagent or resource | Source | Identifier/citation |
|---|---|---|
| **pJLM (modified to remove EGFP)** | Addgene | #19319 |
| **pJLM (modified to remove EGFP)-Halo** | This study | Addgene ID pending |
| **pJLM (modified to remove EGFP)-TFIIB-Halo** | This study | Addgene ID pending |
| **pHA-Kaposin B** | | |
| **pHA-EGFP** | | |
| ***Equipment*** | | |
| **Chemidoc MP Imager** | Bio-Rad | 12003154 |
| **Neon Transfection System** | Invitrogen | MPK5000 (discontinued) |
| **qSonica Ultrasonicator Cuphorn** | qSonica | 431C2 |
| **QuantStudio3 Real-Time PCR** | Applied Biosystems | Available through ThermoFisher |
| **Nikon Eclipse Ti spinning disk confocal microscope** | Nikon | N/A |
| **Qubit 3 Fluorometer** | Invitrogen | Q33216 |
| **Fragment Analyzer** | AATI/Agilent | N/A |
| **CFX Connect thermal cycler** | BioRad | 1855201 |
| **Illumina NovaSeq S1 flowcell** | Illumina | N/A |
| **LSR Fortessa** | BD Bioscience | N/A |

| **Reagent or resource** | **Source** | **Identifier/citation** |
|---|---|---|
| ***Genomes*** | | |
| **GRCh38.p13 human genome assembly and Gencode v43 annotations** | Ensembl | enembl.org |
| **KSHV reference genome** | N/A | Brulois et al. 2012 (18) |
| **KSHV annotations GTF file** | This study | Available upon request |
| ***Deposited Data*** | | |
| ***Software*** | | |
| **Bio-Rad Image Lab v. 6.0.1** | Bio-Rad | SOFT-LIT-170-9690-ILSMAC-V-6-1 |
| **GraphPad Prism v.10.10** | GraphPad Software, Inc. | v.10.10 |
| **Nikon Imaging Systems (NIS) Elements** | Nikon | (v. AR 5.21.03) |
| **FIJI (ImageJ v. 2.0.0-rc-69/1.52p)** | ImageJ | 2.0.0-rc-69/1.52p (22) |
| **HTStream** | https://s4hts.github.io/HTStream/ | N/A- documentation on site |
| **STAR** | https://github.com/alexdobin/STAR | Dobin et al. 2018 (23) |
| **Limma** | https://bioconductor.org/packages/devel/workflows/vignettes/RNAseq123/inst/doc/limmaWorkflow.html | Law et al. (24) |

| Reagent or resource | Source | Identifier/citation |
|---|---|---|
| **FlowJo software** | BD Biosciences; https://www.flowjo.com/ | N/A |
| **BioRender.com** | BioRender; https://www.biorender.com/ | N/A |
| **Spyder** | https://www.spyder-ide.org/ | v. 5.3.3 |
| **Rstudio** | https://www.rstudio.com/categories/rstudio-ide/ | v. 2023.06.2+561 |
| **For details on overall RNA-seq analysis workflow, see UC Davis Bioinformatics course documentation** | https://ucdavis-bioinformatics-training.github.io/2022-August-RNA-Seq-Analysis/ | N/A |

**Supplemental Table 3.** DNA sequences used for cloning and RT-qPCR in this study.

| Function | Sequence | Template | Additional notes |
|---|---|---|---|
| ***Cloning*** | | | |
| **F primer for TFIIB cDNA amplification** | ctgttgtgtcttgttgcggg | cDNA from TRExRTA BCBL-1 | Amplifies TFIIB with some 5' and 3' UTR |
| **R primer for TFIIB cDNA amplification** | aggcccaagctcatcacttc | cDNA from TRExRTA BCBL-1 | Amplifies TFIIB with some 5' and 3' UTR |
| **F primer for TFIIB cloning with XhoI-digested pCDNA/TO-2xStrep (C-terminal)** | caccgcctccCTCGAGtagctgtggtagtttgtccactggg | siRNA resistant TFIIB Twist construct | |
| **F primer for TFIIB cloning with XhoI-digested pCDNA/TO-2xStrep (C-terminal)** | GTGGCGGCCGCTCGAatggcgtctaccagccg | siRNA resistant TFIIB Twist construct | |
| **R primer for TFIIB strep tag SfiI-digested sleeping beauty vector #60511** | ccccaagcttggcctTTAAACGGGCCCcttctcg | TFIIB in pCDNA/TO 2xStrep (C-terminal) | |

| Function | Sequence | Template | Additional notes |
|---|---|---|---|
| **F primer for TFIIB strep tag SfiI-digested sleeping beauty vector #60511** | accccaagctggcctatggcgtctaccagccg | TFIIB in pCDNA/TO 2xStrep (C-terminal) | |
| **F primer for inverse PCR to generate TFIIB(S65A)-2xStrep mutant** | caacaaaagatccagctcgagttgg | TFIIB-2xStrep in #60511 | |
| **F primer for inverse PCR to generate TFIIB(S65E)-2xStrep** | caacaaaagatccagaacgagttggag | TFIIB-2xStrep in #60511 | |
| **R primer for inverse PCR to generate TFIIB (S65A and S65E)-2xStrep** | ctttgtcattgctgaaagttcgcc | TFIIB-2xStrep in #60511 | |
| **F primer for inverse PCR to generate TFIIB(D207A)-2xStrep** | gtccgtggccttgattacaactg | TFIIB-2xStrep in #60511 | |
| **R primer for inverse PCR to generate TFIIB(D207A)-2xStrep** | gtctctagcgctttcaaaataagtttaaagc | TFIIB-2xStrep in #60511 | |
| **F primer for TFIIB-GS-Halo Infusion into Sfi1-digested pLG040. Amplifies TFIIB-GS-Halo homology portion.** | accccaagctggcctatggcgtctaccagccg | si-insentive TFIIB template | 3-way Infusion with Halo amplifcation makes sleeping beauty cloning intermediate for lentivector insertion. Note these include homology regions to add GS linker. |

| Function | Sequence | Template | Additional notes |
|---|---|---|---|
| **R primer for TFIIB_GS_Halo Infusion into Sfi1-digested pLG040. Amplifies TFIIB-GS-Halo homology portion.** | CGATTTCTGCTCCTCCGCTTCCTCCTCCGCTTCCTCCtagctgtggtagtttgtccactg | si-insentive TFIIB template | 3-way Infusion with Halo amplifcation makes sleeping beauty cloning intermediate for lentivector insertion. Note these include homology regions to add GS linker. |
| **F primer for TFIIB-GS-Halo Infusion into Sfi1-digested sleeping beauty vector intermediate. Amplifies TFIIB homology-GS-Halo.** | agctaGGAGGAAGCGGAGGAGGAAGCGGAGGAGCAGAAATCGGTACTGGCTTTCCATTC | Halo template | 3-way Infusion with TFIIB amplifcation makes sleeping beauty cloning intermediate for lentivector insertion. Note these include homology regions to add GS linker. |
| **R primer for TFIIB-GS-Halo Infusion into Sfi1-digested sleeping beauty vector intermediate. Amplifies Halo.** | ccccaagcttggcctCTAGCCGGAAATCTCGAGCG | Halo template | 3-way Infusion with TFIIB amplifcation makes sleeping beauty cloning intermediate for lentivector insertion. Note these include homology regions to add GS linker. |
| **F primer for inserting TFIIB-GS-Halo into EcoRV digested pJLM-ZeoR** | accggTGTCGACGATatggcgtctaccagccg | si-Insensitive TFIIB-GS-Halo from sleeping beauty intermediate | |
| **R primer for inserting TFIIB-GS-Halo into EcoRV digested pJLM-ZeoR** | gaattCTCTAGAGATCTAGCCGGAAATCTCGAGCGTCGAC | si-Insensitive TFIIB-GS-Halo from sleeping beauty intermediate | |
| **F primer for Blasticidin resistance marker replacement into pJLM1-ZeoR** | TCCCCAGGGGGATCCATGCCTTTGTCTCAAGAAGAATCCAC | BlastR variant in #60510 | |

| Function | Sequence | Template | Additional notes |
|---|---|---|---|
| **R primer for Blasticidin resistance marker replacement into pJLM1-ZeoR** | GGTCTTAAAGGTACCCCATAGAGCCCACCGCAT | BlastR variant in #60510 | |
| **R primer for amplifying pJLM1-blast to remove TFIIB to make new Halo only pLJM1 Blast vector.** | catATCGTCGACAccggtagcg | pJLM1-TFIIB-Halo-BlastR | Also adds start codon to Halo protein |
| **F primer for amplifying pJLM1-blast to remove TFIIB to make new Halo only pLJM1 Blast vector.** | GCAGAAATCGGTACTGGCTTTCCATTC | pJLM1-TFIIB-Halo-BlastR | Also adds start codon to Halo protein |
| ***qPCR primers*** | | | |
| **TFIIB forward** | CAGTTCTGATCGGGCAATGAT | Bulk RNA | Performed efficiency correction on these primers |
| **TFIIB reverse** | AGAAATTCGTGATACGGCACAT | Bulk RNA | Performed efficiency correction on these primers |
| **18s forward** | GTGGAGCGATTTGTCTGGTT | Bulk RNA | |
| **18s reverse** | GTGGAGCGATTTGTCTGGTT | Bulk RNA | |
| **K8.1 Promoter forward** | GGGAGAACCATGCCAGACTTTG | Purified bulk DNA | |
| **K8.1 Promoter reverse** | GCATAGGATTAGGAGCGCCAC | Purified bulk DNA | |

| Function | Sequence | Template | Additional notes |
|---|---|---|---|
| **TLDC1 promoter forward** | AGCATGCCCCGACTGCT | Purified bulk DNA | |
| **TLDC1 promoter reverse** | CCCGGAAGGTCAGCGTG | Purified bulk DNA | |

### Supplemental References

1. K. Randolph, U. Hyder, I. D'Orso, KAP1/TRIM28: Transcriptional Activator and/or Repressor of Viral and Cellular Programs? *Frontiers in Cellular and Infection Microbiology* **12** (2022).
2. P. Czerwińska, S. Mazurek, M. Wiznerowicz, The complexity of TRIM28 contribution to cancer. *Journal of Biomedical Science* **24** (2017).
3. C. McCormick, D. Ganem, The Kaposin B Protein of KSHV Activates the p38/MK2 Pathway and Stabilizes Cytokine mRNAs. *Science* **307**, 739-741 (2005).
4. C. A. King, Kaposi's Sarcoma-Associated Herpesvirus Kaposin B Induces Unique Monophosphorylation of STAT3 at Serine 727 and MK2-Mediated Inactivation of the STAT3 Transcriptional Repressor TRIM28. *Journal of Virology* **87**, 8779-8791 (2013).
5. G. Y. Liu *et al.*, Antizyme, a natural ornithine decarboxylase inhibitor, induces apoptosis of haematopoietic cells through mitochondrial membrane depolarization and caspases' cascade. *Apoptosis* **11**, 1773-1788 (2006).
6. P. L. Zhou *et al.*, Perilipin 5 deficiency promotes atherosclerosis progression through accelerating inflammation, apoptosis, and oxidative stress. *Journal of Cellular Biochemistry* **120**, 19107-19123 (2019).
7. J. Feng *et al.*, Perilipin 5 ameliorates high-glucose-induced podocyte injury via Akt/GSK-3β/Nrf2-mediated suppression of apoptosis, oxidative stress, and inflammation. *Biochem Biophys Res Commun* **544**, 22-30 (2021).
8. L. Wang *et al.*, Hypoxia-induced LncRNA DACT3-AS1 upregulates PKM2 to promote metastasis in hepatocellular carcinoma through the HDAC2/FOXA3 pathway. *Experimental & Molecular Medicine* **54**, 848-860 (2022).
9. J. Liang *et al.*, Mitochondrial PKM2 regulates oxidative stress-induced apoptosis by stabilizing Bcl2. *Cell Research* **27**, 329-351 (2017).
10. X. Qu *et al.*, Loss of cancer-associated fibroblast-derived exosomal DACT3-AS1 promotes malignant transformation and ferroptosis-mediated oxaliplatin resistance in gastric cancer. *Drug Resistance Updates* **68**, 100936 (2023).
11. Q. Li *et al.*, Disruption of Robo2-Baiap2 integrated signaling drives cystic disease. *JCI Insight* **4** (2019).
12. S. Qian *et al.*, The role of BCL-2 family proteins in regulating apoptosis and cancer therapy. *Frontiers in Oncology* **12** (2022).
13. Y. Wang *et al.*, ALPK2 acts as tumor promotor in development of bladder cancer through targeting DEPDC1A. *Cell Death & Disease* **12** (2021).
14. C. Meneses *et al.*, The angiotensin-(1–7)/Mas axis reduces myonuclear apoptosis during recovery from angiotensin II-induced skeletal muscle atrophy in mice. *Pflügers Archiv - European Journal of Physiology* **467**, 1975-1984 (2015).
15. L. Zhang, M. Yao, W. Ma, Y. Jiang, W. Wang, MicroRNA-376b-3p targets RGS1 mRNA to inhibit proliferation, metastasis, and apoptosis in osteosarcoma. *Annals of Translational Medicine* **9**, 1652-1652 (2021).
16. N. Labo *et al.*, Heterogeneity and Breadth of Host Antibody Response to KSHV Infection Demonstrated by Systematic Analysis of the KSHV Proteome. *PLoS Pathogens* **10**, e1004046 (2014).

17. E. Hartenian, A. S. Mendez, A. L. Didychuk, S. Khosla, Britt, DNA processing by the Kaposi's sarcoma-associated herpesvirus alkaline exonuclease SOX contributes to viral gene expression and infectious virion production. *Nucleic Acids Research* **51**, 182-197 (2023).
18. K. F. Brulois *et al.*, Construction and Manipulation of a New Kaposi's Sarcoma-Associated Herpesvirus Bacterial Artificial Chromosome Clone. *Journal of Virology* **86**, 9708-9720 (2012).
19. J. Myoung, D. Ganem, Generation of a doxycycline-inducible KSHV producer cell line of endothelial origin: Maintenance of tight latency with efficient reactivation upon induction. *Journal of Virological Methods* **174**, 12-21 (2011).
20. M. Stürzl, D. Gaus, W. G. Dirks, D. Ganem, R. Jochmann, Kaposi's sarcoma-derived cell line SLK is not of endothelial origin, but is a contaminant from a known renal carcinoma cell line. *International Journal of Cancer* **132**, 1954-1958 (2013).
21. H. Nakamura *et al.*, Global Changes in Kaposi's Sarcoma-Associated Virus Gene Expression Patterns following Expression of a Tetracycline-Inducible Rta Transactivator. *Journal of Virology* **77**, 4205-4220 (2003).
22. J. Schindelin *et al.*, Fiji: an open-source platform for biological-image analysis. *Nature Methods* **9**, 676-682 (2012).
23. A. Dobin *et al.*, STAR: ultrafast universal RNA-seq aligner. *Bioinformatics* **29**, 15-21 (2013).
24. C. W. Law *et al.*, RNA-seq analysis is easy as 1-2-3 with limma, Glimma and edgeR. *F1000Research* **5**, 1408 (2018).
